# Cryptic genetic structure and copy-number variation in the ubiquitous forest symbiotic fungus *Cenococcum geophilum*

**DOI:** 10.1101/2021.07.29.454341

**Authors:** Benjamin Dauphin, Maíra de Freitas Pereira, Annegret Kohler, Igor V. Grigoriev, Kerrie Barry, Hyunsoo Na, Mojgan Amirebrahimi, Anna Lipzen, Francis Martin, Martina Peter, Daniel Croll

## Abstract

Ectomycorrhizal (ECM) fungi associated with plants constitute one of the most successful symbiotic interactions in forest ecosystems. ECM support trophic exchanges with host plants and are important factors for the survival and stress resilience of trees. However, ECM clades often harbour morpho-species and cryptic lineages, with weak morphological differentiation. How this relates to intraspecific genome variability and ecological functioning is poorly known. Here, we analysed 16 European isolates of the ascomycete *Cenococcum geophilum*, an extremely ubiquitous forest symbiotic fungus with no known sexual or asexual spore forming structures but with a massively enlarged genome. We carried out whole-genome sequencing to identify single-nucleotide polymorphisms. We found no geographic structure at the European scale but divergent lineages within sampling sites. Evidence for recombination was restricted to specific cryptic lineages. Lineage differentiation was supported by extensive copy-number variation. Finally, we confirmed heterothallism with a single *MAT1* idiomorph per genome. Synteny analyses of the *MAT1* locus revealed substantial rearrangements and a pseudogene of the opposite *MAT1* idiomorph. Our study provides the first evidence for substantial genome-wide structural variation, lineage-specific recombination and low continent-wide genetic differentiation in *C. geophilum.* Our study provides a foundation for targeted analyses of intra-specific functional variation in this major symbiosis.

**Originality-Significance Statement:** We provide the first report on the genetic structure and copy-number variation of the globally ubiquitous and key forest symbiotic fungus *Cenococcum geophilum* using whole-genome sequencing data. We found divergent lineages within sampling sites, while closely related lineages appear over large geographic distances on a continental scale. Even though no sexual spore forming structures have been reported to date, we provide evidence of recombination in a specific lineage suggesting mating activity. Our findings help explain the high genetic diversity occurring within populations and their resilience to changing and adverse environmental conditions. Furthermore, we identify a single *MAT1* idiomorph per genome, confirming heterothallism, and discover that major genomic rearrangements are found in their flanking regions based on chromosomal synteny analysis. Intriguingly, a pseudogene of the opposite functional idiomorph has been characterised in each genome, suggesting a common homothallic ancestor to the species. As *Cenococcum geophilum* is a pivotal mycorrhizal associate of a broad range of trees and shrubs providing nutrition and water supply to their hosts, we highlight and discuss the potential role of the large genome-wide structural variations in environmental selection.

## Introduction

Ectomycorrhizal (ECM) fungi associated with plants constitute one of the most successful symbiotic interactions in forest ecosystems (Martin *et al*., 2016). Abundant ECM partners support the nutrition of many host tree species by facilitating the acquisition of essential nutrients, which is a key determinant of plant survival and productivity under abiotic stress or changing environmental conditions. With exceptional species richness (∼20,000; Martin *et al*., 2016), ECM fungi carry high functional diversity among species that is a key factor of ecosystem resilience (Larsen *et al*., 2016). The environment, through biotic and abiotic factors, plays a major role in shaping ECM communities and their host range. Environmental variation supports ECM species diversity at both small (e.g. host plants) and large spatial scales, and underlies phenotypic plasticity of functional importance for host trees (van der Linde *et al*., 2018). Although intraspecific diversity of important symbiotic traits was highlighted in a handful of studies (Cairney, 1999; Rineau & Courty, 2011), the extent to which such variation is genetically driven and contributes to functional diversity within ECM communities remains largely unknown (Johnson *et al*., 2012).

*Cenococcum geophilum* Fr. (Dothideomycetes, Ascomycota; Spatafora *et al*., 2012) is one of the most ubiquitous and abundant ECM fungi in the world (LoBuglio, 1999). *C. geophilum* colonises a large variety of hosts and occupies a wide ecological niche, from alpine to tropical regions (Trappe, 1962). The species tolerates adverse environmental conditions, as it is commonly found in soils with limited water availability, contaminated by heavy metals, or in sites with stressful climatic conditions such as at the timberline (LoBuglio, 1999; Hasselquist *et al*., 2005; Gonçalves *et al*., 2009; Herzog *et al*., 2013). Increased drought tolerance and resistance to desiccation have been shown for *C. geophilum* mycelia and ECM root-tips as compared to other ECM species (Coleman *et al*., 1989; di Pietro *et al*., 2007; Fernandez *et al*., 2013). These studies also revealed high intraspecific variability and demonstrated that melanin content is one of the key functional traits conferring an advantage under water stress. In nature, *C. geophilum* isolates have shown patterns of local adaptation to serpentine soils with a significant effect of nickel concentrations on fitness-related traits (Gonçalves *et al*., 2009; Bazzicalupo *et al*., 2020). Collectively, these results suggest that this cosmopolitan ECM fungus has undergone selection pressures imposed by various environmental factors.

*Cenococcum geophilum* is typically recognised based on morphology, i.e., a morpho-species. It forms black melanised mycorrhizae with thick, darkly pigmented hyphae spreading from the root tips into the surrounding soil. Although rare in other ECM species (Smith *et al*., 2015), *C. geophilum* produces sclerotia that act as a “sclerotial bank” in the soil, which are resistant propagules that are melanised and can withstand desiccation for several decades and sequester carbon as dead sclerotia for centuries (Watanabe *et al*., 2007; Obase *et al*., 2014, 2018). Although being common, the biology and life cycle remains poorly understood. Phylogenetic analyses have revealed multiple divergent *C. geophilum* lineages on small spatial scales, even at the level of soil core samples (Douhan and Rizzo, 2005; Obase *et al*., 2016). Combined with the high genetic diversity within populations (ECM root tips and sclerotia), this suggests that *C. geophilum* has undergone cryptic speciation (Obase *et al*., 2017; Vélez *et al*., 2021).

Our understanding of *C. geophilum* reproduction is limited. Whilst no spore-producing structures were observed, indicating strictly asexual reproduction through vegetative mycelial propagation, including germination of sclerotia, population genetic studies suggested recombination events that would have involved sexual mating (LoBuglio and Taylor, 2002; Douhan *et al*., 2007). However, mating may well be restricted to certain environmental conditions or to a few specific lineages with distinct gene pathways involved in sexuality, as suggested in arbuscular mycorrhizal fungi (Riley and Corradi, 2013). This is supported by phylogenetic studies that showed little gene flow within *C. geophilum* populations. Subtle geographic structure with long-distance disjunction suggests (Obase *et al*., 2016) a complex alternation of sexual and asexual reproduction over space and time (Obase *et al*., 2017). Besides further sampling efforts, analyses at a whole genome level are essential to determine the prevalence of outcrossing and to better understand patterns of cryptic speciation.

The genome of *C. geophilum,* the only known ECM member within Dothideomycetes, is uniquely large for this class (Peter *et al*., 2016). A high-quality reference genome (strain 1.58) revealed that about 75% of the genome is composed of transposable elements (TEs) and this high proportion is unevenly distributed in the genome, with TE-rich and TE-poor regions (de Freitas Pereira *et al*., 2018). Its closest sequenced relatives within Gloniaceae, Mytilinidiales, the saprotrophic species *Glonium stellatum* and *Lepidopterella palustris,* for example, showed a fourfold smaller genome size (Peter *et al*., 2016). TE proliferation seems to be a hallmark for the transition from saprotrophy to symbiosis and in fact, *C. geophilum* is the ECM species with the highest proportion of TEs known to date among mycorrhizal fungi with one of the largest genomes, i.e., 178 Mbp; Miyauchi *et al*., 2020). Nevertheless, the gene content (14,748) is similar to the average number of genes found in fungi (Miyauchi *et al*., 2020). Intraspecific genome size variation suggests important structural rearrangements or gene duplications and deletions (Bourne *et al*., 2014). Investigation of the reference genome sequence highlighted the presence of sex-related genes and an intact *MAT1* locus containing one of the two idiomorphs (here *MAT1-1*) typical for a heterothallic bipolar mating system of ascomycetes (Peter *et al*., 2016). The mating-type locus carrying the opposite idiomorph, i.e., *MAT1-2,* has not been analysed in *C. geophilum* so far.

In this study, we used whole-genome sequencing to determine the population genetic structure of 16 *C. geophilum* isolates across Europe and test for recombination events. We reconstructed genome-wide phylogenetic relationships within the genus and characterised gene duplication and deletion events to investigate structural patterns of genetic diversity. Then, we analysed patterns of selection at the gene and genome level using nucleotide diversity, Tajima’s *D*, and the ratio of non-synonymous to synonymous polymorphisms. Lastly, we analysed the genetic architecture of the mating type locus and its recent sequence evolution.

## Results

### Read mapping, variant calling, and polymorphism filtering

Illumina sequencing of 16 *C. geophilum* strains resulted in 348.9 million read pairs (Fig. 1A; Table S1). Of those, 293.6 million read pairs (84.16%) were successfully mapped on the reference strain 1.58. The highest and lowest number of raw read pairs was available for the strains 1.101 and 1.094, respectively, and the highest (95.54%) and lowest (64.37%) overall alignment with the reference strain was found in strains 1.061 and 1.096, respectively (Table S1). An average of 11.3 million read pairs were used for mapping (Table S1). We used the Genome Analysis Toolkit (GATK) to detect genetic variants from the 16 sequenced strains and found, after filtering, a total of 345,358 single nucleotide polymorphisms (SNPs), including 284,479 bi-allelic SNPs, of which 67,186 were synonymous variant sites based on the SnpEff analysis.

**Figure 1:**
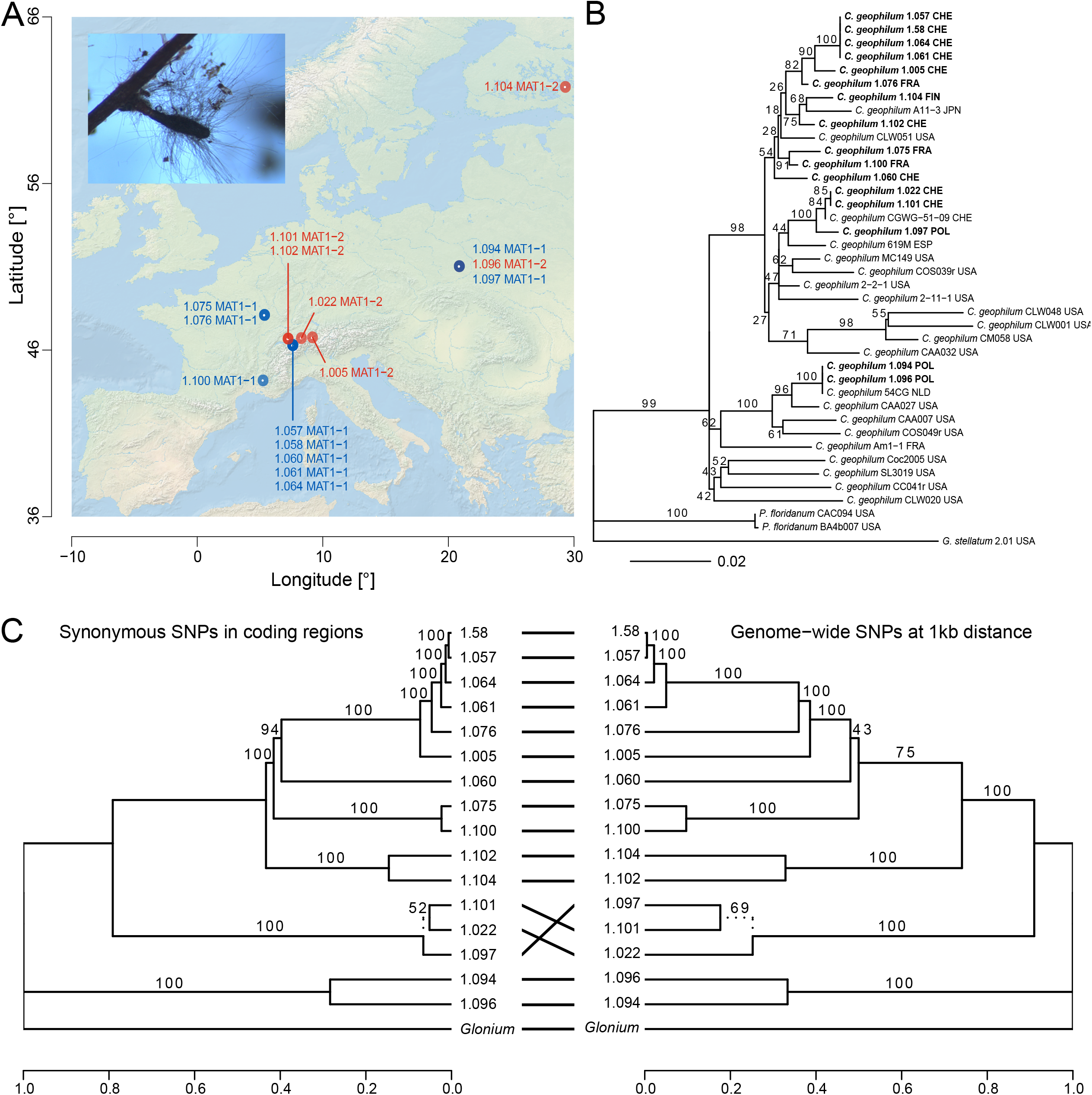
Study range, phylogenetic relationships, and genomic relatedness of the *Cenococcum geophilum* strains sampled. **A**) Sampling sites of the *C. geophilum* isolates with an illustration of the ecto-mycorrhiza morphology (Photo: M. Peter). Geographic distribution of MAT loci is given for each sampled location. **B**) Phylogenetic relationships within the genus *Cenococcum* based on two nuclear loci, i.e., ITS and GAPDH. One outgroup species, i.e., *Glonium stellatum,* was used to root the phylogenetic tree. The scale bar shows the substitution rate. Tips in bold are the 16 samples analysed in this study using whole-genome sequencing data. **C**) Genomic relatedness among the *C. geophilum* sampled, including the reference genome (strain 1.58). Whole-genome SNP data capture both repetitive and coding regions while synonymous SNPs only reflect neutral divergence in coding regions. Topologies are rooted with *Glonium stellatum* as outgroup. Branch supports represent bootstrap values from the maximum likelihood tree reconstruction. Strain codes are described in Table S1.

### Phylogenetic tree reconstructions

We inferred phylogenetic trees based on the GAPDH and ITS loci using maximum-likelihood (ML) analyses to identify the main lineages of *C. geophilum* strains. Our two-locus nuclear dataset contained alignment lengths of 597 and 949 bp for GAPDH and ITS, respectively, with a total of 1,546 aligned bp for the 40 strains (including outgroups; Table S2). The GAPDH locus had the most variable sites. The inferred GAPDH and ITS ML tree topologies revealed several incongruencies, but the concatenated partitions generated a fairly well-resolved phylogenetic tree, with a majority of branches with BS values > 80; Figs. 1B, S1–2). With minor exceptions, we found strong support for several monophyletic clades but those were not congruent with the geographical origin of the samples (Fig. 1B). Overall, the clades that emerged from the sequenced European strains were depicted in the worldwide clades 5 and 6 described by Obase et al. (2016).

To improve resolution, we used genome-wide data and found highly congruent tree topologies compared to the GAPDH and ITS phylogeny. The phylogenomic tree was robust to the use of two near neutral datasets of either synonymous SNPs or randomly-selected genome-wide SNPs (Fig. 1C). Both genome-based ML trees were well-resolved with most of branches being strongly supported (BS > 90; Fig. 1C). This indicates that neutral divergence in the coding regions matches the phylogenetic relationships given by the divergence in non-coding or coding regions.

### Recombination events and genetic structure

We analysed patterns of recombination using a Neighbour-Net network in SplitsTree and found evidence for significant events among the genomes of *C. geophilum* strains (Fig. 2A). We identified two major genetic groups connected by two strains (1.005 and 1.076). The Φ-test supported recombination across the genome when all *C. geophilum* were included (*P* < 0.001; Table S3). However, applying the _Φ_ test to only the cluster not containing the reference genome (non-ref. cluster) yielded no significant evidence for recombination (*P* = 0.641), suggesting one clonal and one recombinant genetic group.

**Figure 2:**
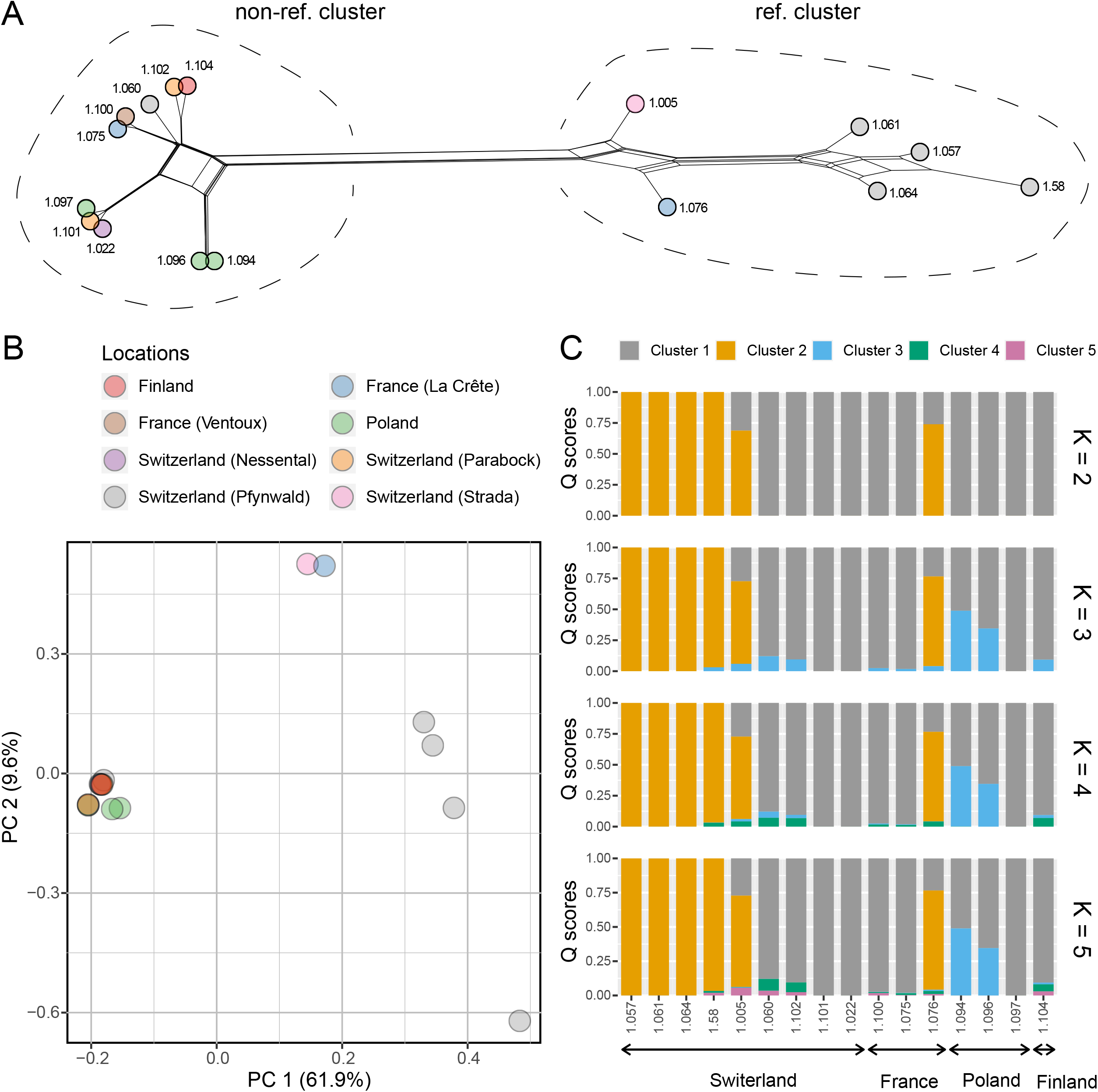
Recombination analysis and population genetic differentiation among the *Cenococcum geophilum* strains sampled. **A**) Network-based phylogenetic reconstruction supporting recombination events in one of the two main lineages. **B**) Principal component (PC) analysis showing the genomic variation without clear congruence with geography. **C**) Ancestry inferences for *K* = 2–5 indicating some admixed genotypes between the major genetic groups. Strain codes are described in Table S1. Colours refer to the geographic origin and genetic group of each strain for A–B and C, respectively.

As for the phylogenetic analyses, we found no clear geographic structure based on either principal components (PC) or Bayesian clustering analysis. For example, isolates from the Pfynwald (CHE) sampling site (1.057, 1.58, 1.060, 1.061, and 1.064) clustered into the two major groups, reporting a new case of highly divergent lineages present in the same sampling site (Fig. 2A). In addition, isolates from the Blizyn (POL) sampling site (1.094, 1.096, and 1.097) also showed divergent lineages, but these were only represented in the clonal group with both mating types (Fig. 2A). No fungal lineages exclusive to certain host tree species were observed in this dataset. The reference genome 1.58 appeared at the edge of the recombinant genetic group. The first axes of the PCA captured 61.9% and 9.6% of the genetic variance (PC1 and PC2, respectively; Fig. 2B). In total, the first three axes explained 76.8% of the overall genotypic variation (Fig. S3). Consistent with the Neighbour-Net network, PC1 and PC2 separated three to four genetic groups with substantial differentiation between the Pfynwald strains. The intermediate genetic group consisting of the 1.005 and 1.076 strains also clustered together independently. In addition, Bayesian clustering analyses were congruent between replicates (Fig. S4) and we found the optimal number of clusters to be *K* = 2 (Fig. 2C). At *K* = 2, two major groups were clearly identified and the intermediate group (strains 1.005 and 1.076), displayed in the PCA and network analysis, belonged to the Pfynwald group containing the sequenced strain 1.58. At *K* = 3, the intermediate group still exhibited patterns of admixture between the two major groups, and two Polish genotypes (strains 1.094 and 1.096 belonging to the phylogenetic Clade 6 in Obase *et al*., 2016) indicate contributions from a third genetic group. At higher *K* values, no significant changes were observed in genotype ancestry memberships.

### Detection of copy-number variations

We conducted copy-number variation (CNV) analyses comparing read depth of *C. geophilum* strain sequencing sets against the reference genome. We improved the robustness of CNV calls by removing any CNV calls in regions where self-alignments of strain 1.58 reads against the 1.58 reference genome produced candidate CNVs. Overall, we found that gene duplications occurred ∼3.5 fold more frequently than deletions (Fig. 3A). This predominance of gene duplications was only detected at the highest rates of overlap between genes and CNV regions, i.e., >95% similarity. In total, we identified 217 and 72 gene duplications and deletions, respectively, using a threshold of >50% similarity. By inferring the overall tree topology of based on CNV genotypes, we found patterns consistent with the outcomes of the genetic clustering and phylogenetic analyses (Fig. 3B). First, five strains (1.022, 1.094, 1.096, 1.097, and 1.101) formed a monophyletic clade that was paraphyletic in either the two-locus or genome-based phylogenies (Figs. 1C, 3B). Second, we identified a monophyletic clade with six strains (1.060, 1.075, 1.100, 1.102, 1.104, and 1.58) that constituted a paraphyletic clade in the previous analyses. Furthermore, we tested whether the distance between genes affected by CNVs and the closest TEs are smaller than those for all genes as indicated for other species (e.g. Hartmann and Croll, 2017). Interestingly, CNV genes were at a slightly larger distance compared to the background (*P* = 0.028, 719 and 448 median base pairs, respectively; Fig. 4).

**Figure 3:**
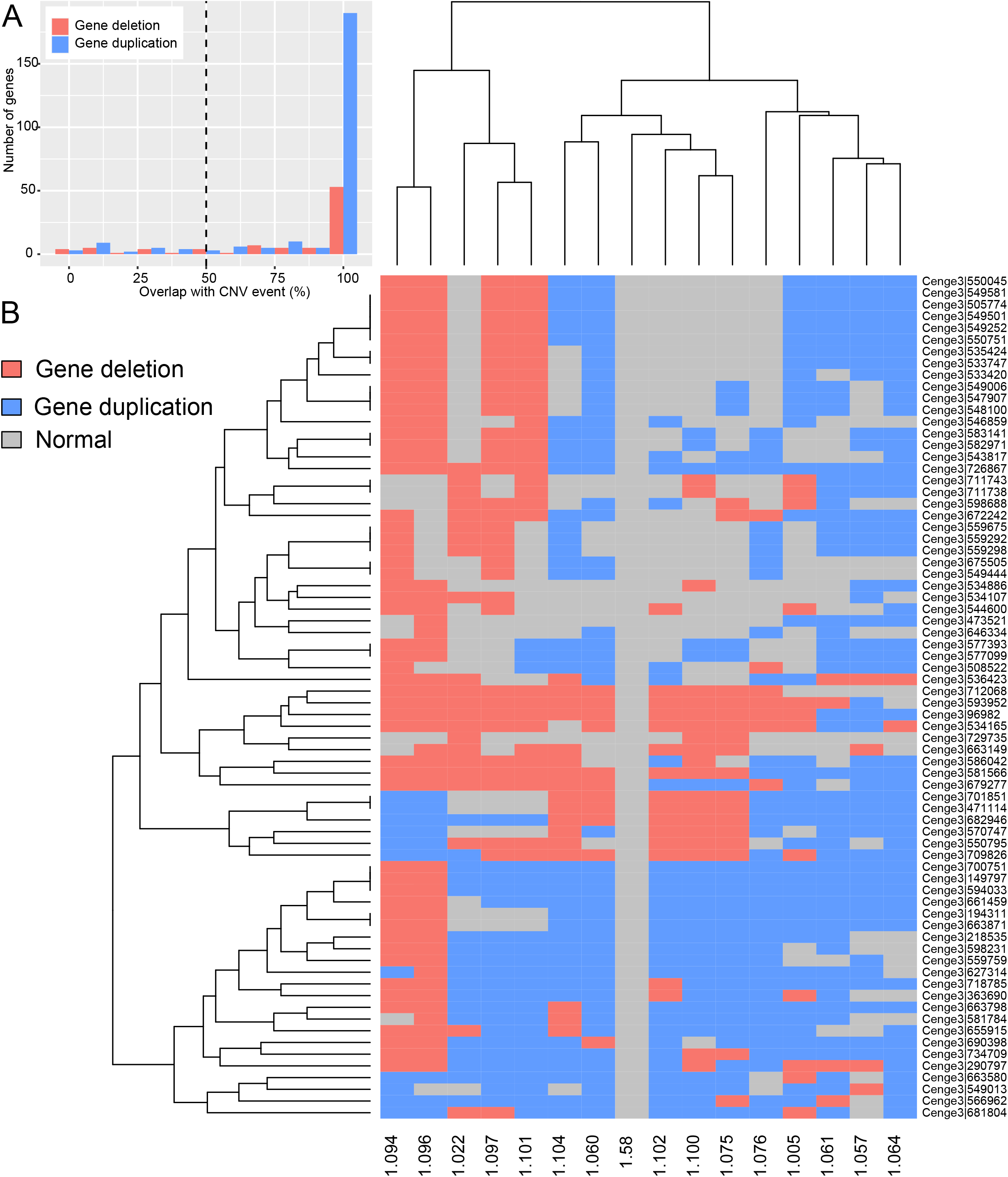
Analysis of copy-number variation (CNV) regions among the sampled *Cenococcum geophilum* strains. **A**) Gene content including CNV with the percentage of length covered by deletion or duplication >50%. Deletions refer to any gene locus showing at least one deletion in a strain while duplications include any gene locus showing only duplication or normal copy-numbers without deletion. **B**) Gene comparison affected by deletion or duplication among *C. geophilum* strains. Tree topologies on the left and top show the grouping according to gene CNVs. The y-axis shows gene identifiers of CNV-affected genes. Note that the strain 1.58 displays only single-copy genes because the CNV calls were filtered to contain only those genes that were confirmed as single-copy from the reference genome. Strain codes are described in Table S1.

**Figure 4:**
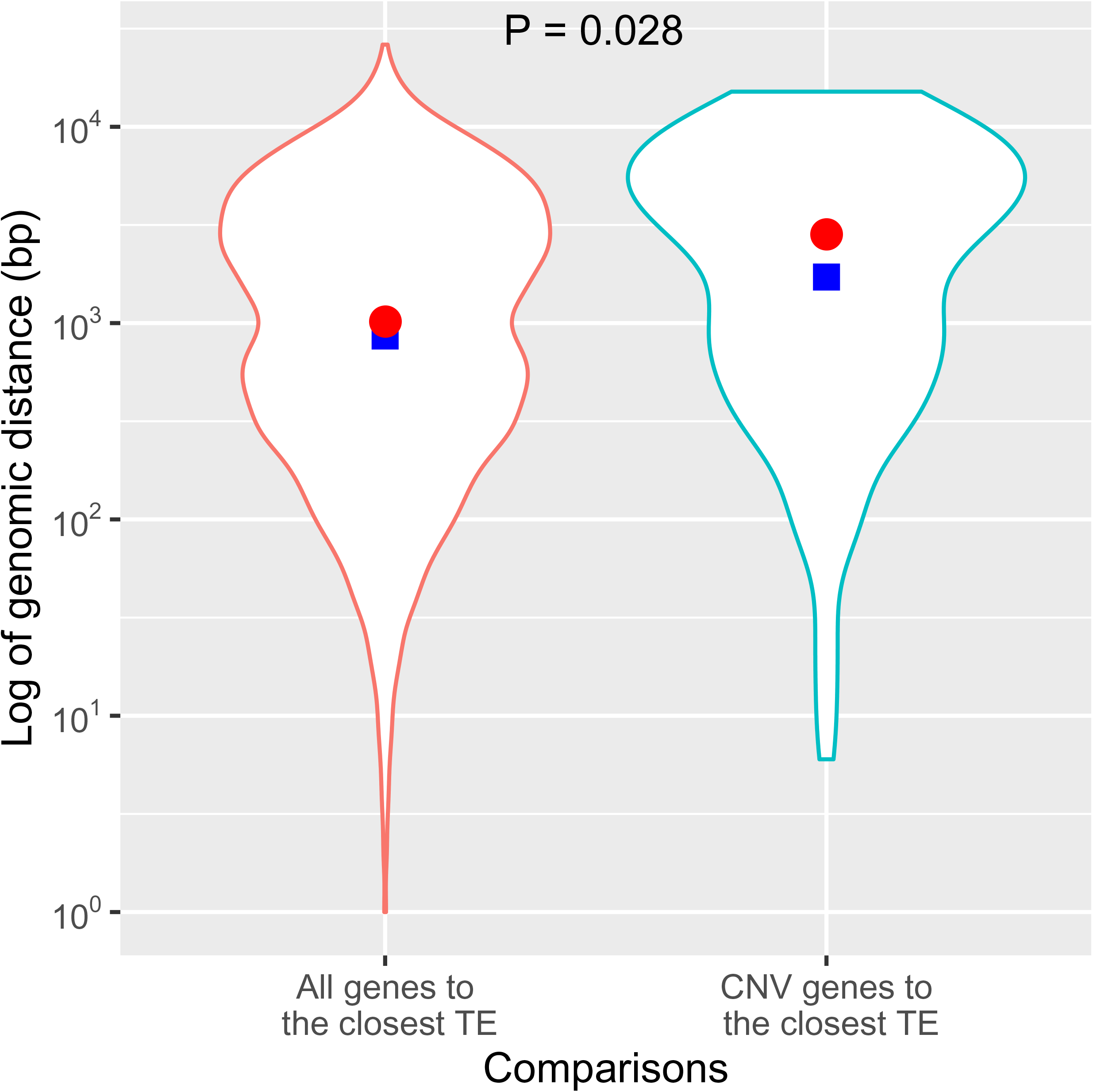
Distribution of genomic distances between all genes and their closest transposable elements (TEs), and between copy-number variation (CNV) genes and their closest TEs. The significance between the median values was calculated using the Wilcoxon test. The red dots and blue squares represent mean and median values, respectively.

### Within-species genetic diversity and signatures of selection

We assessed within-species genetic diversity across the 16 *C. geophilum* genomes and identified genomic regions with elevated diversity rates along the two longest scaffolds, i.e., #1 and #2; Fig. 5A,C, on which the density of coding sequences and transposable elements (TEs) were negatively correlated (Fig. 5B,D). At the gene level, we estimated the nucleotide diversity per site (π) to be 0.000–0.020, 0.000–0.028, and 0.000–0.013 for the “all”, “ref. cluster”, and “non-ref. cluster” datasets, respectively (Fig. 6A). The genome-wide average π value was 3.31×10^-3^, 1.31×10^-3^, and 1.01×10^-3^, respectively (Fig. 6A). These results support that the ref. cluster is genetically more diverse than the non-ref. cluster, although the sample size is nearly double in the latter. At the gene-level, we calculated Tajima’s *D* values ranging from −1.90–2.86, −0.87–2.32, and −1.97–2.42 for the “all”, “ref. cluster”, and “non-ref. cluster” dataset, respectively (Fig. 6B). At the genome-wide level, the average Tajima’s *D* estimates were 0.14 (ref. cluster) and 0.31 (non-ref. cluster), with a value of 1.64 for the “all” dataset (Fig. 6B). This strongly skewed Tajima’s *D* distribution when all strains are included towards positive values indicates strong population substructure (Fig. 6B). However, the “ref. cluster” dataset showed a non-normal distribution of Tajima’s *D* with an excess of negative values, which is indicative of a population expansion. The non-ref. cluster showed a distribution largely centred around 0 under neutrality and constant population size. A slight deviation towards positive values suggests a historic decrease in population size or some population substructure. The distribution of the *p*N/*p*S ratio assessed from the all dataset suggests purifying selection (Fig. 6C).

**Figure 5:**
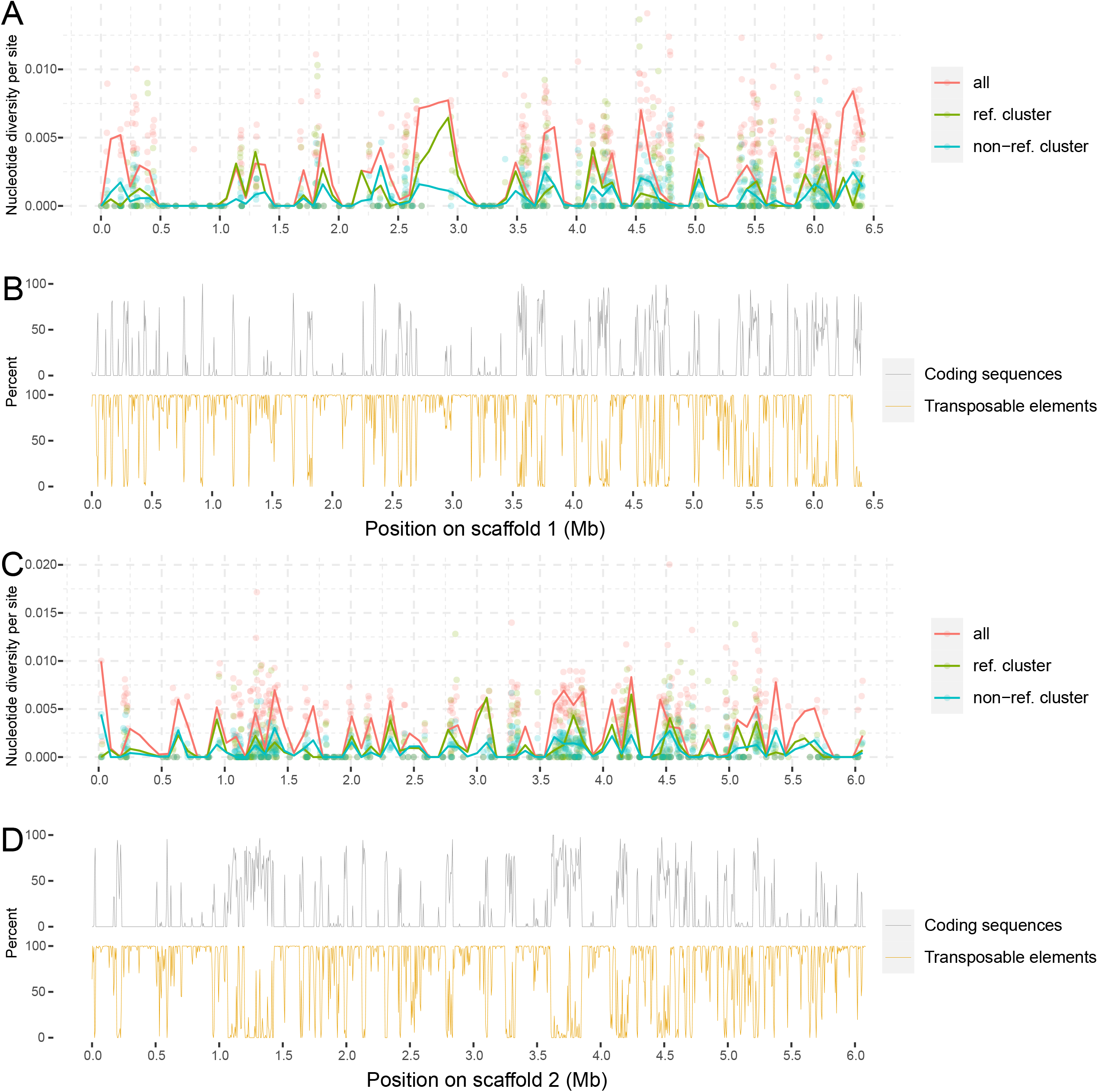
Genetic diversity analysis and coding sequence variation along scaffolds #1 and #2 in *Cenococcum geophilum* strains used in this study. **A**) and **C**) Nucleotide diversity per site is given for each gene identified in the 1.58 genome assembly. **B**) and **D**) Density of gene content and transposable element is presented. Results are shown for the complete dataset (“all”) and subsets (“ref. cluster” and “non-ref. cluster”) of *Cenococcum geophilum* strains used in this study (see section Experimental Procedures).

**Figure 6:**
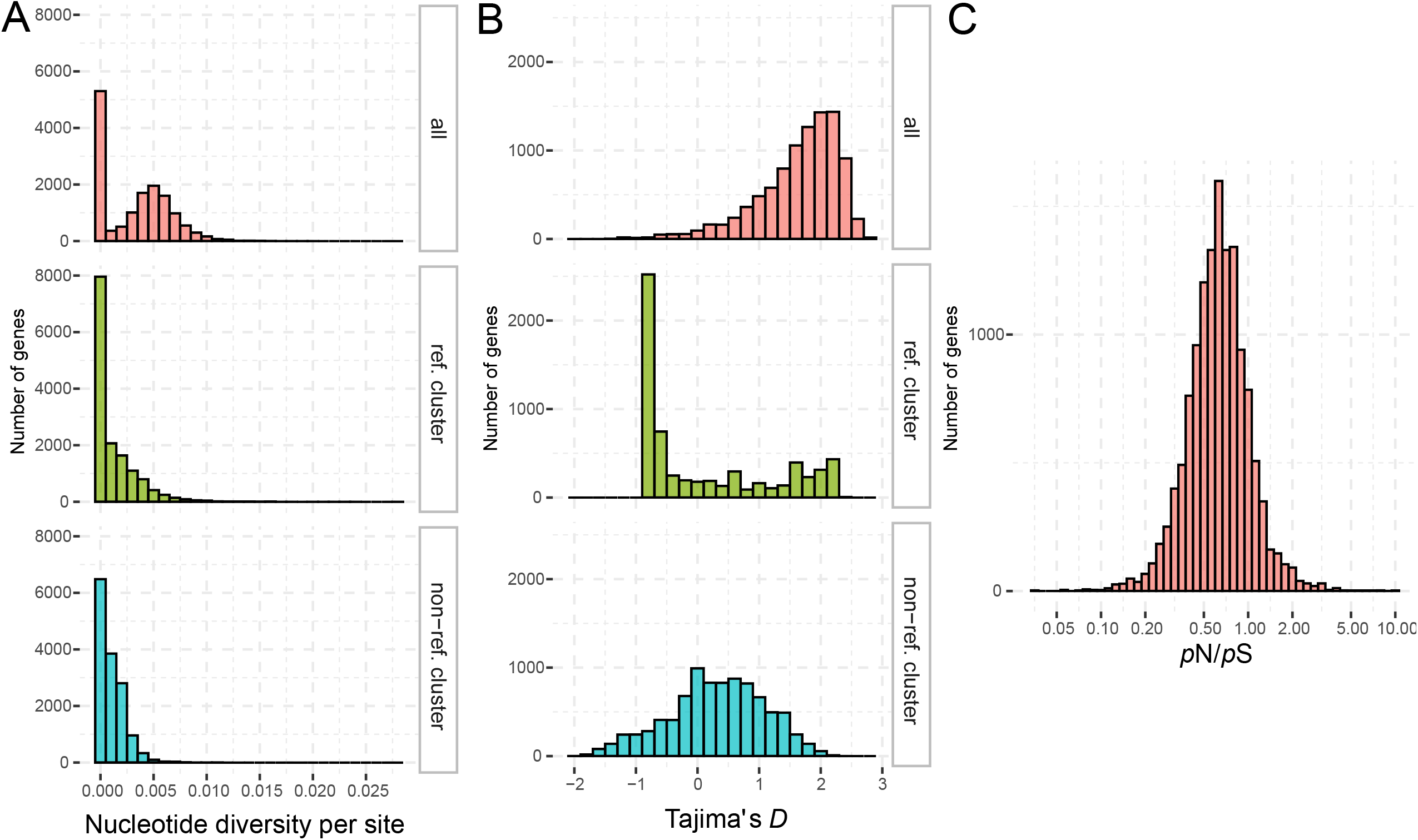
Within-species genetic diversity and signatures of selection. **A**) Distribution of nucleotide diversity per site for each gene. **B**) Distribution of gene-specific Tajima’s *D*. **C**) Distribution of the gene-specific ratio of non-synonymous to synonymous polymorphisms (*p*N/*p*S). Results are shown for the complete dataset (“all”) and subsets (“ref. cluster” and “non-ref. cluster”) of *Cenococcum geophilum* strains used in this study (see section Experimental Procedures). Note that the reference cluster is genetically more diverse than the non-ref. cluster.

Then, we characterised the top-candidate genes putatively under selection by identifying those with the lowest Tajima’s *D* estimates in either the “ref. cluster” or “non-ref. cluster” (see Experimental procedures). Based on functional annotation, we found that those top-candidate genes differed to some extent between the ref. and non-ref. clusters (Table 2). In the recombining “ref. cluster”, increased numbers of genes were detected that are putatively involved in self/non-self recognition containing HET, NACHT, WD40- and Ankyrin-repeat domains, as well as genes that are members of multigene-families that are expanded in *C. geophilum* or unique in this species and its closest relatives *G. stellatum* and *L. palustris* as compared to other Dothideomycetes (Peter *et al*., 2016). In the “non-ref. cluster”, we found increased number of genes involved in the secondary metabolism (SM; e.g. Lofgren *et al*., 2021) including two genes found in SM clusters for prenyltransferase synthesis (DMATs; Khaldi *et al*., 2010). In both ref. and non-ref. clusters, we found one gene coding for a carbohydrate-active enzyme (CAZyme; Cantarel *et al*., 2009). In the “ref. cluster”, it codes for a secreted β-galactosidase (Zeiner *et al*., 2016), a member of the glycoside hydrolyse family 35 (GH35) implicated in hemicellulose degradation (van den Brink and de Vries, 2011) that is contracted in the *C. geophilum* genome as compared to other Dothideomycetes. In the “non-ref. cluster”, it is a member of the lytic polysaccharide monooxigenases (LPMO) family A11 that are almost exclusively present in ascomycetes and the one functionally characterised so far involved in chitin degradation (Hemsworth *et al*., 2014). Other genes found in both groups encode for proteins involved in information storage and processing as well as cellular processes and signalling (Table S4). All these genes were expressed in free-living mycelium and/or functioning ECM root-tips, but most of them not differentially regulated in these tissues and if so, then mostly down-regulated in ECMs (Table S4).

**Table 1:** Details of the Cenococcum geophilum individuals analysed in this study.

| Strain | *MAT1* locus | Isolate origin | Code | Site | Forest | Isolation year | Latitude [° N] | Longitude [° E] |
| --- | --- | --- | --- | --- | --- | --- | --- | --- |
| 1.104 | *MAT1-2* | Finland | FIN | Punkaharju | *Pinus sylvestris* | 20.08.08 | 61.800000 | 29.316667 |
| 1.076 | *MAT1-1* | France | FRA | La Crête | *Fagus sylvatica* | 1999 | 48.111111 | 5.356944 |
| 1.075 | *MAT1-1* | France | FRA | La Crête | *Fagus sylvatica* | 1999 | 48.111111 | 5.356944 |
| 1.100 | *MAT1-1* | France | FRA | Ventoux | Mixed | 30.07.09 | 44.179497 | 5.241850 |
| 1.094 | *MAT1-1* | Poland | POL | Blizyn | Mixed | 01.03.09 | 51.037222 | 20.869083 |
| 1.096 | *MAT1-2* | Poland | POL | Blizyn | Mixed | 13.08.09 | 51.037111 | 20.868694 |
| 1.097 | *MAT1-1* | Poland | POL | Blizyn | Mixed | 13.08.09 | 51.037222 | 20.870250 |
| 1.022 | *MAT1-2* | Switzerland | CHE | Nessental | *Picea abies* | 13.06.80 | 46.722500 | 8.296667 |
| 1.101 | *MAT1-2* | Switzerland | CHE | Parabock | *Picea abies* | 15.05.08 | 46.690960 | 7.236940 |
| 1.102 | *MAT1-2* | Switzerland | CHE | Parabock | *Picea abies* | 10.10.08 | 46.690960 | 7.236940 |
| 1.057 | *MAT1-1* | Switzerland | CHE | Pfynwald | *Pinus sylvestris* | 23.04.08 | 46.301389 | 7.611944 |
| 1.58 | *MAT1-1* | Switzerland | CHE | Pfynwald | *Pinus sylvestris* | 30.04.08 | 46.301389 | 7.611944 |
| 1.061 | *MAT1-1* | Switzerland | CHE | Pfynwald | *Pinus sylvestris* | 30.04.08 | 46.301389 | 7.611944 |
| 1.064 | *MAT1-1* | Switzerland | CHE | Pfynwald | *Pinus sylvestris* | 04.09.08 | 46.301389 | 7.611944 |
| 1.060 | *MAT1-1* | Switzerland | CHE | Pfynwald | *Pinus sylvestris* | 30.04.08 | 46.301389 | 7.611944 |
| 1.005 | *MAT1-2* | Switzerland | CHE | Strada | *Picea abies* | 03.11.77 | 46.767500 | 9.178889 |

**Table 2:**
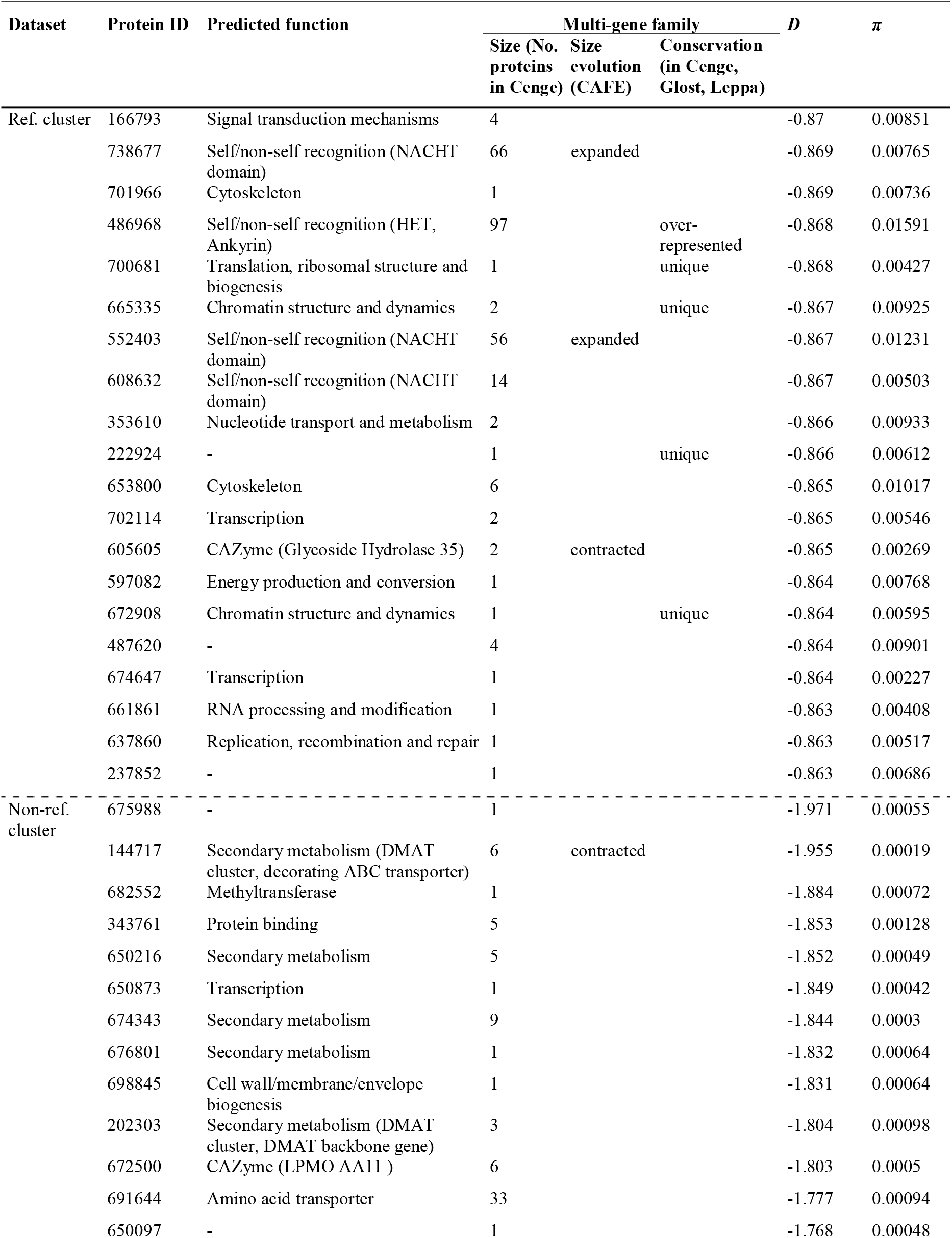
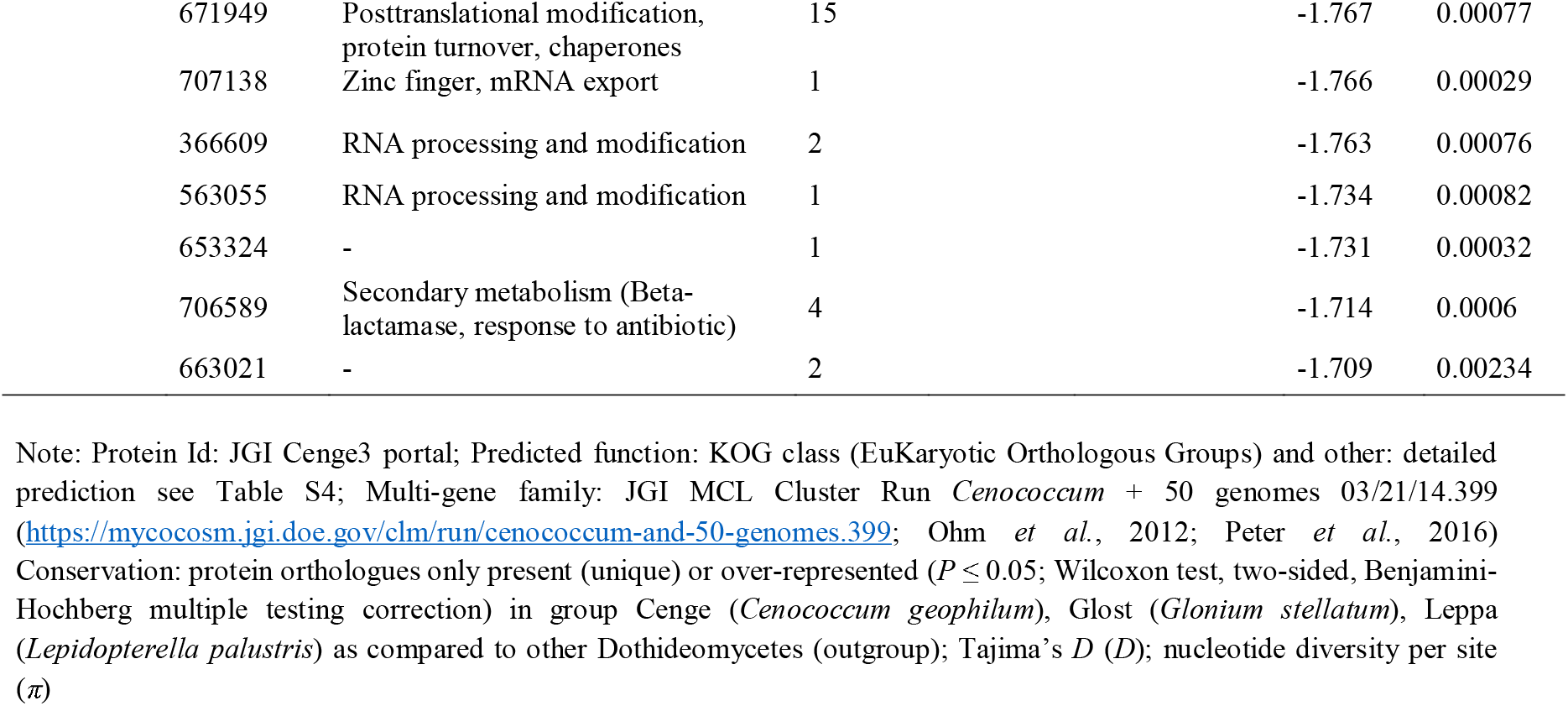
Characteristics of the 20 top-candidate genes putatively under selection in the ref. and non-ref. cluster based on Tajima’s D statistics. Estimation of the nucleotide diversity per site is shown for each candidate gene.

| Dataset | Protein ID | Predicted function | Multi-gene family | | | $D$ | $\pi$ |
| --- | --- | --- | --- | --- | --- | --- | --- |
|  |  |  | Size (No. proteins in Cenge) | Size evolution (CAFE) | Conservation (in Cenge, Glost, Leppa) |  |  |
| Ref. cluster | 166793 | Signal transduction mechanisms | 4 |  |  | -0.87 | 0.00851 |
|  | 738677 | Self/non-self recognition (NACHT domain) | 66 | expanded |  | -0.869 | 0.00765 |
|  | 701966 | Cytoskeleton | 1 |  |  | -0.869 | 0.00736 |
|  | 486968 | Self/non-self recognition (HET, Ankyrin) | 97 |  | over-represented | -0.868 | 0.01591 |
|  | 700681 | Translation, ribosomal structure and biogenesis | 1 |  | unique | -0.868 | 0.00427 |
|  | 665335 | Chromatin structure and dynamics | 2 |  | unique | -0.867 | 0.00925 |
|  | 552403 | Self/non-self recognition (NACHT domain) | 56 | expanded |  | -0.867 | 0.01231 |
|  | 608632 | Self/non-self recognition (NACHT domain) | 14 |  |  | -0.867 | 0.00503 |
|  | 353610 | Nucleotide transport and metabolism | 2 |  |  | -0.866 | 0.00933 |
|  | 222924 | - | 1 |  | unique | -0.866 | 0.00612 |
|  | 653800 | Cytoskeleton | 6 |  |  | -0.865 | 0.01017 |
|  | 702114 | Transcription | 2 |  |  | -0.865 | 0.00546 |
|  | 605605 | CAZyme (Glycoside Hydrolase 35) | 2 | contracted |  | -0.865 | 0.00269 |
|  | 597082 | Energy production and conversion | 1 |  |  | -0.864 | 0.00768 |
|  | 672908 | Chromatin structure and dynamics | 1 |  | unique | -0.864 | 0.00595 |
|  | 487620 | - | 4 |  |  | -0.864 | 0.00901 |
|  | 674647 | Transcription | 1 |  |  | -0.864 | 0.00227 |
|  | 661861 | RNA processing and modification | 1 |  |  | -0.863 | 0.00408 |
|  | 637860 | Replication, recombination and repair | 1 |  |  | -0.863 | 0.00517 |
|  | 237852 | - | 1 |  |  | -0.863 | 0.00686 |
| Non-ref. cluster | 675988 | - | 1 |  |  | -1.971 | 0.00055 |
|  | 144717 | Secondary metabolism (DMAT cluster, decorating ABC transporter) | 6 | contracted |  | -1.955 | 0.00019 |
|  | 682552 | Methyltransferase | 1 |  |  | -1.884 | 0.00072 |
|  | 343761 | Protein binding | 5 |  |  | -1.853 | 0.00128 |
|  | 650216 | Secondary metabolism | 5 |  |  | -1.852 | 0.00049 |
|  | 650873 | Transcription | 1 |  |  | -1.849 | 0.00042 |
|  | 674343 | Secondary metabolism | 9 |  |  | -1.844 | 0.0003 |
|  | 676801 | Secondary metabolism | 1 |  |  | -1.832 | 0.00064 |
|  | 698845 | Cell wall/membrane/envelope biogenesis | 1 |  |  | -1.831 | 0.00064 |
|  | 202303 | Secondary metabolism (DMAT cluster, DMAT backbone gene) | 3 |  |  | -1.804 | 0.00098 |
|  | 672500 | CAZyme (LPMO AA11 ) | 6 |  |  | -1.803 | 0.0005 |
|  | 691644 | Amino acid transporter | 33 |  |  | -1.777 | 0.00094 |
|  | 650097 | - | 1 |  |  | -1.768 | 0.00048 |

|  |  |  |  |  |
| --- | --- | --- | --- | --- |
| 671949 | Posttranslational modification,<br>protein turnover, chaperones | 15 | -1.767 | 0.00077 |
| 707138 | Zinc finger, mRNA export | 1 | -1.766 | 0.00029 |
| 366609 | RNA processing and modification | 2 | -1.763 | 0.00076 |
| 563055 | RNA processing and modification | 1 | -1.734 | 0.00082 |
| 653324 | - | 1 | -1.731 | 0.00032 |
| 706589 | Secondary metabolism (Beta-<br>lactamase, response to antibiotic) | 4 | -1.714 | 0.0006 |
| 663021 | - | 2 | -1.709 | 0.00234 |

Note: Protein Id: JGI Cenge3 portal; Predicted function: KOG class (EuKaryotic Orthologous Groups) and other: detailed prediction see Table S4; Multi-gene family: JGI MCL Cluster Run *Cenococcum* + 50 genomes 03/21/14.399 (https://mycocosm.jgi.doe.gov/clm/run/cenococcum-and-50-genomes.399; Ohm *et al.*, 2012; Peter *et al.*, 2016) Conservation: protein orthologues only present (unique) or over-represented ( $P \leq 0.05$ ; Wilcoxon test, two-sided, Benjamini-Hochberg multiple testing correction) in group Cenge (*Cenococcum geophilum*), Glost (*Glonium stellatum*), Leppa (*Lepidopterella palustris*) as compared to other Dothideomycetes (outgroup); Tajima's $D$ ( $D$ ); nucleotide diversity per site ( $\pi$ )

### Chromosomal synteny analysis of the MAT1-1 and MAT1-2 idiomorphs

Our *de novo* draft genome assemblies of other strains than 1.58 generated 151,48 million base pairs (Mbp) on average, with a mean number of 26,963 contigs per assembly and a mean largest contig size of 165,938 bp. The value range for N50 and L50 were 5,720–24,382 and 2,247–4,759, respectively (Table S5). The GC content varied from 33 to 40%, with an average of 37%. Mismatches were lowest and highest with 61 and 260 per 100 kbp for the 1.101 and 1.094 strains, respectively.

We identified two idiomorphs at the *MAT1* locus in *C. geophilum* isolates and found genetic rearrangements in its flanking coding and non-coding regions based on the synteny analysis (Fig. 7). A single functional *MAT1* idiomorph was detected in each strain, supporting the heterothallism of the species (Figs. 1, 7; Table 1). In both *MAT1* loci types, we found a pseudogene of the opposite *MAT1* idiomorph close to the functional one showing conserved strand orientation as compared to strains carrying the functional opposite idiomorphs with unchanged relative position of their open reading frames (ORF). The same was observed for the closely related *G. stellatum (MAT1-2*: Glost2|444166), whereas *L. palustris*, a homothallic species and also member of the Mytilinidiales, carries both functional *MAT1* idiomorphs with conserved orientation but rearranged position (*MAT1-1*: Leppa1|291743; *MAT1-2*: Leppa1|291655). The *MAT1-1* and *MAT1-2* idiomorphs were surrounded by a conserved gene encoding a MFS transporter (555485) and a gene coding for a protein (72340) of unknown function, both of which showing evidence for an insertion polymorphism generating a non-syntenic region among the *MAT1-1* and *MAT1-2* strains. Furthermore, the flanking *APN2* gene (686539) coding for a putative DNA lyase, and the conserved protein TEX2 (634502) coding for a putative PH domain-containing integral membrane protein were also present in the *MAT1* loci of the closely related *G. stellatum* and *L. palustris* and are often found adjacent to *MAT1* idiomorphs in ascomycetes (Debuchy *et al*., 2014; Wang *et al*., 2016; APN2: Glost2|386243, Leppa1|414878; TEX2: Glost2|444168, Leppa1|322867). We further found a gene (555074) encoding a cysteine-type peptidase and a protein of unknown function (72410).

**Figure 7:**
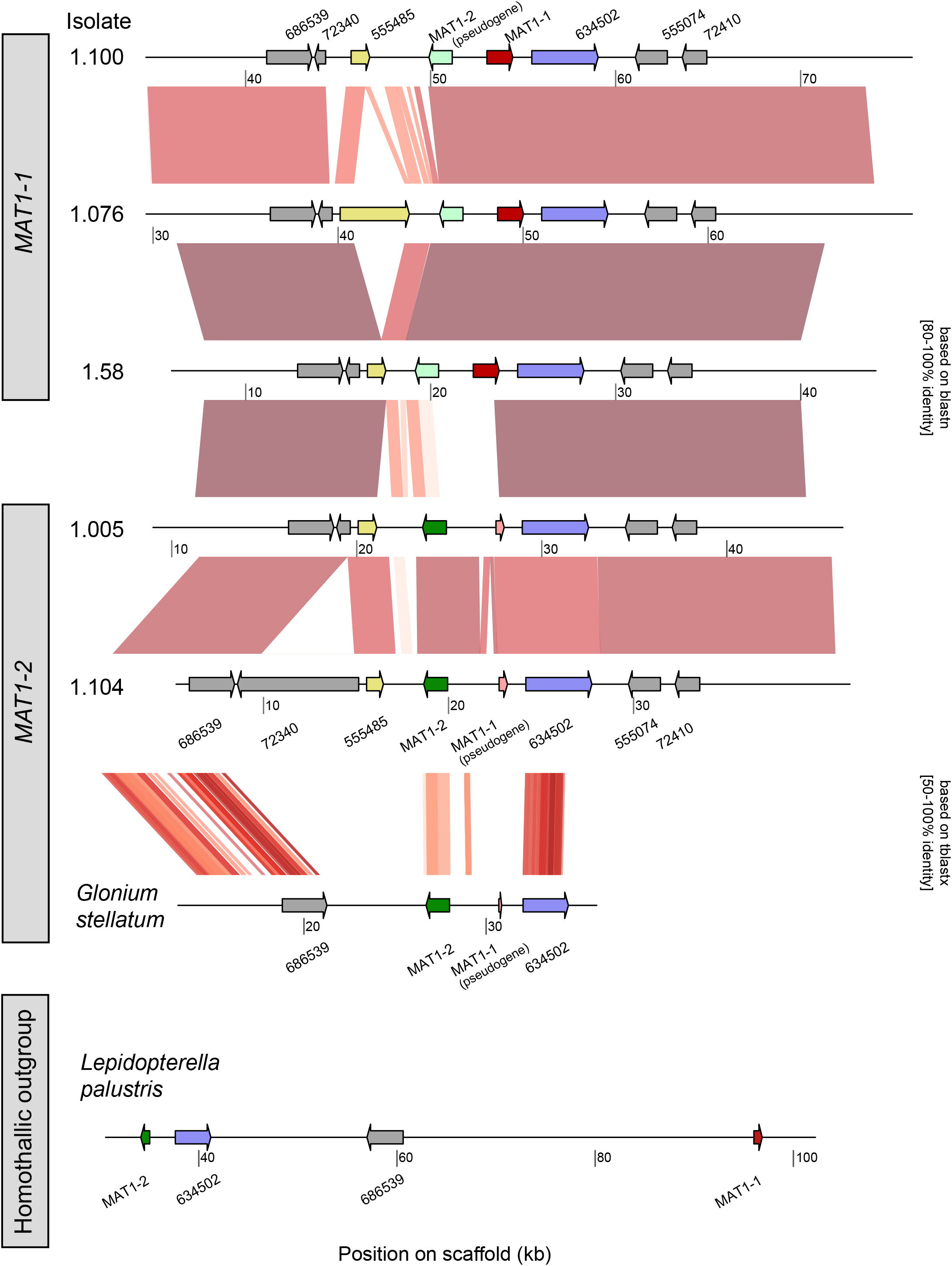
Chromosomal synteny analyses of *MAT1-1* and *MAT1-2* loci. Intraspecific analyses were performed with the reference genome strain (1.58) and interspecific comparisons were carried out between *G. stellatum*, *L. palustris* and *C. geophilum* strains. Colour gradients (white to red) correspond to sequence similarities ranging from 50–100%, with a minimum bit-score of 100.

## Discussion

### Cryptic population genetic structure and evidence for recombination

Our genomic analyses of European *C. geophilum* strains, which showed that divergent lineages co-occur in single sampling sites without major geographic structuring, support previous reports from North America that were based on a restricted number of markers (Douhan and Rizzo, 2005; Vélez *et al*., 2021). Interestingly, our network analysis indicated that the two main genetic groups had possibly different levels of recombination, with most of the sampled strains being clustered into the clonal group with little gene flow. It is unclear whether this restricted gene flow is due to incompatible mating systems or clonal proliferation. Since neither sexual nor asexual spore forming structures have been reported for *C. geophilum*, this suggests a vegetative mycelial propagation in the soil with limited migration ability of the species. In the recombinant group, cryptic sexual events must have occurred at some point in the near or distant past. In case of a asexually reproducing population, i.e., mating incompatibility, low genetic similarities are expected between strains sampled in the same site, which could explain the observed pattern within the Pfynwald site, where one strain (1.060) clearly differs and did not group in the recombinant cluster (Fig. 2A). Our synteny analyses of the *MAT1* loci showed important rearrangements in close vicinity to the *MAT1* genes, but whether this affects mating compatibility is unclear. Apart from mating incompatibility or sexual versus vegetative propagation as well as isolation by geographic distance, other evolutionary processes could explain the current genetic structure, such as demographic history, past fluctuations in effective population size (e.g. population expansion or contraction), or processes of adaptation to environmental conditions during glacial and interglacial periods, as reported in other ECM fungi (Geml *et al*., 2006; Sánchez-Ramírez *et al*., 2015).

Alternatively, recent adaptation of *C. geophilum* to new hosts or environmental conditions may have led to the emergence of cryptic lineages with strong genetic structure over short geographical distances. While *C. geophilum* is known to have a broad host tree spectrum, some lineages may nevertheless show host preferences, as reported in other ECM species (Rochet *et al*., 2011; Lofgren *et al*., 2021). In this respect, the tree-fungus interaction might have shaped differentiation and genetic isolation of these cryptic lineages, in combination with recent selection. The present data do not support such a mechanism because no lineage grouping according to the dominant host tree species was observed. Also, plant-fungus synthesis experiments showed that functional ectomycorrhizas are formed with diverse tree species, both broadleaves and conifers, for the reference strain 1.58 but also for others (e.g. de Freitas Pereira *et al*., 2018). This shows that the symbiosis can be established with diverse hosts, even if there are possibly host preferences under natural conditions. To shed light on host specificity as a potential driver of genetic structuring within and among sites, detailed information on *C. geophilum* populations associated with roots of different host tree species is needed. Regarding the environment, adaptation to soil characteristics may prevent wide distribution, foster differentiation through geographical isolation and, ultimately, lead to independent and clonal lineages. Note that edaphic specialisation can be coupled with, or even supplanted by, the competition avoidance strategy of ECM species within soil communities (Mujic *et al*., 2016). Clearly, our dataset with 16 strains is limited and further population genomic studies, with more isolates per populations, are required to be conclusive about the prevalence of sexual reproduction under varying ecological conditions throughout the species range.

### Massive gene duplications and functional regions under positive selection

Our CNV analysis revealed broad genomic alterations among *C. geophilum* lineages with about 3.5-fold more gene duplications than deletions. Interestingly, the genome-wide pattern of CNV genotypes is congruent with the tree topology from phylogenetic reconstructions suggesting that CNVs have accumulated gradually since the origin of the species. However, the genomic distances between the CNV genes and their closest TEs were in the same range as those for all genes. This is unexpected since previous studies showed that genes affected by CNVs are often close to TEs due to their mutagenic potential (e.g. Hartmann and Croll, 2017). A reason for this finding might be the high abundance of TEs throughout the *C. geophilum* genome. Although de Freitas *et al*. (2018) found islands of TE-rich and TE-poor regions, these were rather small and scattered throughout the scaffolds so that most genes have TEs in close vicinity. With about 75% of its genome comprising TEs, *C. geophilum* exhibits the highest proportion of transposons and retrotransposons found in ECM fungi (18– 75%; Miyauchi *et al*., 2020) and other Dothideomycetes (1–75%; Peter *et al*., 2016). Our findings corroborate the recent proliferation of species-specific TEs and its evolutionary significance in affecting functional rearrangements of the genome (Daboussi and Capy, 2003; Muszewska *et al*., 2019). However, other mechanisms such as aneuploidy, chromosomal duplication, unequal crossing over may also be responsible for these gene duplications (Bourne *et al*., 2014). Such high intraspecific genomic diversity has lately been observed for other long-considered asexual fungi, such as the ancient arbuscular mycorrhizal glomeromycetes, and TE activity was one of the explanatory mechanism (Chen *et al*., 2018). Likewise, genomic studies indicated the presence of cryptic sex in these species, which, as in our case, might explain the high adaptability of these fungi to a wide range of hosts and habitats and likely has functional significance in host-environment interactions (Chen *et al*., 2018; Reinhardt *et al*., 2021).

As in another ascomycete ECM fungus, *Tuber melanosporum* (Payen *et al*., 2015), we found a large proportion of genes that exhibited an excess of low frequency polymorphisms relative to neutral expectation (Tajima’s *D* < –1, non-ref. cluster; Fig. 6B). Candidate genes identified as putatively under selection need to be interpreted with caution given our limited sample size and heterogeneous genetic structure. Nevertheless, a large number of genes identified in the recombining ref. cluster are putatively implicated in self/non-self recognition containing HET, NACHT, WD40- and Ankyrin-repeat domains and are members of expanded gene families. Such HET domain proteins were also identified as putative candidates under selection for the outcrossing ECM ascomycete *T. melanosporum* in a similar analysis of six isolates covering its geographical distribution (Payen *et al*., 2015). Fungal genomes contain a large number of genes encoding HET domains of unknown function, but resolved functions often relate to vegetative incompatibility (e.g. conspecific recognition; Paoletti, 2016), hetero-specific, or even inter-kingdom recognition as seen for characterised NOD-like receptors (Dyrka *et al*., 2014). These genes were not up- or down-regulated in ECM root tips as compared with free-living mycelium in a synthesis experiment (Peter *et al*., 2016) and, hence, are likely not involved in specific symbiotic functions in *C. geophilum.* The repeat domains have been shown to evolve under positive diversifying selection (Dyrka et al 2014) as were some groups of HET domain containing proteins that displayed higher allelic diversity and a restricted phylogenetic distribution (Paoletti et al 2016). In the more clonal non-ref. cluster, for which Tajima’s *D* values were generally more negative and centred around 0, we found increased number of candidate genes putatively under selection, which are involved in the secondary metabolism. This matches findings from fungal pathogens (e.g. Hartmann *et al*., 2018). Interestingly, clearer signatures of positive selection were found for an outcrossing fungal pathogen species as compared to a more clonally reproducing generalist fungal pathogen (Derbyshire, 2020).

### Genomic rearrangements of the MAT1 locus

The bipolar mating-type (*MAT*) system governs sexual reproduction in ascomycetes and has a key role in establishing cell identity and determining the sexual cycle. Two fungal partners can mate only if they carry different *MAT* idiomorphs. Based on our chromosomal synteny analysis, we showed that both mating-types exist in *C. geophilum* isolates with the *MAT1-1* or the *MAT1-2* idiomorph. In fact, after screening the sixteen genome sequences, a single *MAT1* idiomorph was consistently found in each strain, thus confirming the heterothallism of the species by excluding autogamy. Since the first in-depth genomic analysis of the *MAT1* locus of the *C. geophilum* reference genome (strain 1.58; Peter *et al*., 2016), the genomic architecture of the intact *MAT1* locus was assumed to be stable and no intraspecific reorganisation was reported in the species. Here, we highlight that both *MAT1-1* and *MAT1-2* loci have undergone important rearrangements with gene expansions in surrounding regions. Surprisingly, we identified a *MAT1-1* pseudogene close to the *MAT1-2* idiomorph, i.e., about 3 kbp, and reciprocally, a *MAT1-2* pseudogene close to the *MAT1-1* idiomorph, i.e., about 2 kbp. This raises the question of whether a homothallic ancestor may have existed and been the precursor of independent heterothallic lineages. Interestingly, this *MAT1-1* pseudogene, also found in *G. stellatum*, is relatively similar to the functional *MAT1-1* idiomorph characterised in the homothallic outgroup *L. palustris* (Fig. 7), with the same orientation and an equivalent gene length. Moreover, the gene sequence of the *MAT1-2* idiomorph appears to be enlarged in both *C. geophilum* and *G. stellatum* compared to *L. palustris*, indicating important protein modification. Overall, the genetic makeup of the *MAT1-1* and *MAT1-2* pseudogenes most likely originates from a common ancestor shared with *L. palustris*, from which distinct mating types evolved. Such mating shifts, from heterothallism to homothallism, were highlighted in the *Neurospora* genus, with up to six independent transition events assessed from 43 taxa (Nygren *et al*., 2011). Distinct genetic mechanisms enabled these transitions (e.g. translocation and unequal crossover) with the mediation of specific transposons that reshaped the genetic architecture of the *MAT* locus (Gioti *et al*., 2012). The high mobility of genetic material in the surrounding *MAT* locus is intriguing and points to the role of TE activities and associated silencing mechanisms such as RIP in shaping the gene architecture and diversity that may orchestrate sexual reproduction in *C. geophilum* (Serrato-Capuchina and Matute, 2018). In several fungal species, TEs were described within or in flanking regions of the *MAT* locus, e.g. in *Blastomyces dermatitidis* (Li *et al*., 2013), *Ceratocystis* sp. (Simpson *et al*., 2018), or *T. melanosporum* (Martin *et al*., 2010). Together, these studies support the idea that TEs are frequently inserted at *MAT* loci. They are therefore key evolutionary forces for genome dynamics likely important not only in pathogenic fungal host and environment interactions as earlier stated (Raffaele and Kamoun, 2012), but also for symbiotic interactions, as indicated by increased proliferation of these elements in ECM fungal species, above all in *C. geophilum* (Miyauchi *et al*., 2020). Further population genomic studies including genomic architecture and prevalence of *MAT* loci within populations are needed to fully understand the success of this widespread and abundant ECM species and its role in forest ecosystems.

### Conclusions

Our study provides one of the first intraspecific genomic comparison of an ECM fungal species sampled on a regional and continental scale. Using whole-genome sequencing from 16 European isolates of the ubiquitous forest symbiotic fungus *Cenococcum geophilum*, we found divergent lineages in geographically restricted located sampling sites and no strong geographic structuring. Our genome-wide polymorphism analyses support subdivisions within the species and suggest two distinct main genetic groups, one more clonal and one recombinant. The phylogeny of lineages and groupings are largely supported by the abundant copy-number variations found among genomes. Our gene diversity analyses indicate higher genetic diversity in the recombinant group even though the clonal cluster contained almost double as many strains. Although these results need to be interpreted with caution, because of the limited number of strains studied and an uneven sampling, the top-candidate genes putatively under positive selection based on Tajima’s *D* statistics differed between the two groups. More genes from lineage-specific, expanded gene families involved in self/non-self recognition were found in the recombinant cluster, whereas genes involved in the secondary metabolism detected in the more clonal cluster. Lastly, we confirmed the heterothallism of *C. geophilum* based on the chromosomal synteny analysis of the *MAT1-1* and *MAT1-2* idiomorphs and uncovered important genetic rearrangements in their surrounding coding and non-coding regions for both, strains carrying the same or opposite *MAT1* idiomorphs. Collectively, these findings demonstrate complex genome architectures in this important ECM symbiont that maybe due to cryptic sex and/or TE-based mechanisms, as similarly suggested for the long-believed asexual arbuscular mycorrhizal fungi. Future studies will elucidate the role that this genetic diversity plays in symbiotic function and the potential for adaptation to changing environments.

## Experimental Procedures

### Study range and selection of isolates

In this study, we selected 16 *Cenococcum geophilum* isolates to cover as broad as possible the European range of the species and the divergent phylogenetic lineages previously reported (Obase *et al*., 2016). All strains were isolated from sclerotia collected below diverse host tree species, i.e., broadleaves and conifers, to maximise intra-specific genetic diversity in close and distant sampling sites. After screening, both mating types were included to investigate sexual reproduction from natural populations (Fig. 1; Table 1). Pure cultures of *C. geophilum* were grown in liquid culture containing a Cg-Medium, i.e., a modified Melin Norkran’s (MMN) medium comprising casein (Kerner *et al*., 2012), and incubated at room temperature in the dark. We cultivated isolates for a month and harvested mycelia that were then pulverised in liquid nitrogen for storage at –80°C until processing.

### DNA extraction and barcoding

Genomic DNA was extracted using the PowerMax Soil DNA extraction kit (MoBio, now DNeasy PowerMax Soil kit of Qiagen) according to the manufacturer’s instructions. DNA was quantified by spectrophotometry (Qubit, Thermo Fisher Scientific, Waltham, USA), and the quality was inspected on agarose gel (1.5%). A minimum of 1 µg of DNA was sent to the US Department of Energy Joint Genome Institute (JGI) for genome resequencing.

To identify the lineage of each isolate, we amplified and sequenced two phylogenetically informative loci at the intraspecific level, the glyceraldehyde 3-phosphate dehydrogenase (GAPDH) gene and the internal transcribed spacer (ITS) region (Obase *et al*., 2016), using Sanger dideoxy sequencing. The optimised polymerase chain reaction (PCR) programs for ITS and the GAPDH gene were conducted following Obase *et al*. (2014). For the GAPDH gene, we used universal primers, as reported in Berbee *et al*. (1999), and for ITS, we used ITS1F and ITS4 (White *et al*., 1990; Gardes and Bruns, 1993). PCR products were cleaned up using the illustra™ ExoProStar™ enzymatic PCR and sequence reaction clean kit according to the manufacturer’s instructions (Merck KGaA, Darmstadt, Germany). Cycle sequencing was performed using the BigDye™ Terminator v3.1 Cycle Sequencing kit and products were purified with the BigDye XTerminator™ purification kit (Thermo Fisher Scientific) before to analyse them on an Applied Biosystems ABI3130 genetic analyser. Loci were sequenced with both forward and reverse primers to ensure unambiguous chromatogram profiles and all sequences were deposited to NCBI (MZ570241-MZ570257; MZ577329-MZ577345).

### Genome sequencing and reference genome sequences

Plate-based DNA library preparation for Illumina sequencing was performed on the Sciclone NGS robotic liquid handling system (PerkinElmer, Waltham, USA) using KAPA Biosystems library preparation kit (Roche, Basel, Switzerland). For each sample, 200 ng of DNA was sheared to 600 base pairs (bp) using a Covaris LE220 focused-ultrasonicator (Woburn, Massachusetts, USA). The sheared DNA fragments were size selected by double-SPRI and then the selected fragments were end-repaired, A-tailed, and ligated with Illumina compatible sequencing adaptors from IDT containing a unique molecular index barcode for each sample library. The prepared library was quantified using KAPA Biosystem’s next-generation sequencing library qPCR kit and run on a Roche LightCycler 480 real-time PCR instrument. The quantified library was then multiplexed with other libraries, and the pool of libraries was then prepared for sequencing on the Illumina HiSeq sequencing platform utilising a TruSeq paired-end cluster kit, v4, and Illumina’s cBot instrument to generate a clustered flow cell for sequencing (San Diego, California, USA). Sequencing of the flow cell was performed on the Illumina HiSeq 2500 sequencer using HiSeq TruSeq SBS sequencing kits, v4, following a 2×100 indexed run recipe at the Joint Genome Institute (JGI; Walnut Creek, California, USA).

We accessed the *C. geophilum* raw reads data from the NCBI Short Read Archive (Table S1) and used the *Glonium stellatum* genome as an outgroup species (Peter *et al*., 2016; https://genome.jgi.doe.gov/Glost2) in downstream analyses. The *C. geophilum* reference genome sequence and gene annotation data (filtered models) for the strain 1.58 were downloaded from the JGI (Peter *et al*., 2016; https://genome.jgi.doe.gov/Cenge3). The sequenced strains included the reference genome strain 1.58 to control for the quality of SNP calling (Peter *et al*., 2016).

### Read mapping, variant calling, and polymorphism filtering

We downloaded sequencing data using fastq-dump, as implemented in the SRA-Toolkit v. 2.9.0 (http://ncbi.github.io/sra-tools/), and inspected read quality with FastQC v. 0.11 (Andrews, 2017). We performed quality trimming of raw reads with Trimmomatic v. 0.36 (Bolger *et al*., 2014) based on the following quality filters: ILLUMINACLIP:TruSeq3-PE.fa:2:30:10, SLIDINGWINDOW:5:10, LEADING:10, TRAILING:10, and MINLEN:50. Trimmed reads were mapped on the reference genome 1.58 using Bowtie v. 2.3 with the - very-sensitive and -local settings (Langmead *et al*., 2009). The resulting BAM files were sorted by position and indexed with SAMtools v. 1.6 (Li *et al*., 2009).

We applied the Genome Analysis Toolkit (GATK) v. 3.7 and used the HaplotypeCaller to perform the SNP and indel discovery (McKenna *et al*., 2010). Individual SNP calls were merged and jointly genotyped using the GATK GenotypeGVCFs tool. Then, we filtered the joint Variant Call Format (VCF) file using hard filtering parameters according to GATK best practices recommendations (Van der Auwera *et al*., 2013). The following thresholds were applied for filtering: AN ≥ 12, i.e., at least 12 isolates had a callable genotype, QUAL ≥ 5,000, QD ≥ 5.0, MQ ≥ 20.0, ReadPosRankSum, MQRankSum and BaseQRankSum each comprised between –2.0 and 2.0. We further classified SNP loci as synonymous or non-synonymous variant sites using SnpEff v. 4.3p (Cingolani *et al*., 2012).

### Phylogenetic tree reconstructions

To infer the phylogenetic relationships of the 16 *C. geophilum* strains within the seven main clades described in Obase *et al*. (2016), we assembled a broader dataset (GAPDH and ITS) and reconstructed a two-loci phylogenetic tree based on maximum likelihood (ML) analysis. The additional strains (22 *C. geophilum*, one *Pseudocenococcum floridanum*, and one *G. stellatum*) covered most of the phylogenetic diversity and intercontinental distribution of the species. This includes important disjunctions in North America and potential cryptic taxa in European Clades 5 and 6 (Obase *et al*., 2016), which were investigated using a coalescent-based species delimitation approach (Zhang *et al*., 2013). For each locus, we aligned sequences using MAFFT v. 7 with default parameters (Katoh *et al*., 2019) and excluded ambiguous regions using Gblocks v.0.91b with the less stringent settings (Castresana, 2000). For each locus, we implemented the optimal substitution model, based on AICc scores calculated with the program ModelTest-NG v. 0.1.7 (Darriba *et al*., 2019; Table S2). We ran ML tree searches with a randomised maximum parsimony starting tree and gamma-distributed site rates using RAxML-NG v. 1.0.0 (Kozlov *et al*., 2019; Edler *et al*., 2021). We performed 20 runs and set 1,000 bootstrap replicates for each run, computed on the best-scoring ML tree with a bootstrap random-number seed. We fixed *G. stellatum* as the outgroup to root phylograms.

At the genome-wide level, we generated two SNP datasets to infer independent phylogenetic tree reconstructions. First, from the overall SNP data genotyped in all 17 strains (16 *C. geophilum* and one *G. stellatum*), we randomly selected a single SNP per 1 kilo base-pair (kbp) of the reference genome sequence using VCFtools v. 0.1.15 (Danecek *et al*., 2011). Second, from the overall SNP dataset, we retained only synonymous variants identified in coding regions (see below SnpEff). We used each SNP dataset for an independent ML tree reconstruction with RAxML-NG (Kozlov *et al*., 2019; Edler *et al*., 2021). We ran the best ML tree by analysing 20 different starting trees and performed 1,000 bootstrap replicates on the best-scoring ML tree with a bootstrap random-number seed. We implemented a General Time Reversible (GTR) substitution model and allowed for rate heterogeneity (GTR+G).

### Recombination events and genetic structure

To gain insights into recombination events and genetic structure, we generated a SNP dataset for intraspecific genetic analyses of *C. geophilum* using the following criteria: a single SNP randomly selected per 1 kbp of the reference genome and genotyped in a least 90% of the strains, i.e., genotyped in at least 15 out of 16 strains. We used SplitsTree v. 4.14.6 to build up a Neighbour-Net network from all *C. geophilum* strains (Huson and Bryant, 2006), and uncorrected p distances calculated from the SNP matrix, which is the proportion of positions at which two sequences differ, to generate the network. The network was drawn based on equal angle splits. Tests of recombination within genetic groups of strains were carried out using the Φ test implemented in SplitsTree (Bruen *et al*., 2006). The test identifies evidence of recombination based on phylogenetic compatibilities of neighbouring SNPs in the matrix. We ran the test with the default settings of a 100-characters window size on the SNP matrix. Three genotype datasets were analysed to assess lineage-specific patterns; comprising all strains (referred as “all”), a reference cluster which contained the reference genome 1.58 (referred as “ref. cluster” including 1.005, 1.057, 1.061, 1.064, 1.076, and 1.58), and a non-reference cluster which did not contain the reference genome (referred as “non-ref. cluster” including 1.022, 1.060, 1.075, 1.094, 1.096, 1.097, 1.100, 1.101, 1.102, and 1.104).

We conducted Bayesian unsupervised genetic clustering using STRUCTURE v. 2.3.4 (Pritchard *et al*., 2000). We used an admixture model with correlated frequencies and no prior regarding the population of origin (Falush *et al*., 2003). The *K* parameter was tested for values ranging from 1 to 5, with 10 repetitions for each tested *K* value. We used 50,000 iterations as a burn-in period and 100,000 iterations per run for the Monte Carlo Markov Chain (MCMC) replicates. We visually inspected the convergence of parameters and diagnosed the optimal *K* value based on the stabilisation of –Ln(*K*) values. We performed a principal component analysis (PCA) to explore genotypic variability based on the SNP matrix using the R package SNPRelate v. 1.20.1 (Zheng *et al*., 2012).

### Detection of copy-number variations

We identified segmental deletions and duplications in sequenced isolates compared to the reference genome 1.58 using CNVcaller (Wang *et al*., 2017). CNVcaller uses normalised read-depth of short read alignments to identify high-confidence copy-number variation (CNV) regions. The window size for the CNV calling was set to 500 bp according to the recommendations based on sequencing coverage. The lower and upper GC content limits were set to 0.2 and 0.7, respectively, and the upper limit of gap content was set to 0.5. We required a minimum of Pearson’s correlation coefficient between two adjacent non-overlapping windows to be >0.5, as recommended for less than 30 samples. We also specified that only a single individual displayed a potential gain/loss event to define a candidate CNV window (-h 1 and -f 0.05) given the heterogeneity in genotypes and small sample size. CNV regions were further filtered based on the normalised read depth. Deletions and duplications were retained if the normalised read depth was <0.4 and >1.8, respectively. CNV regions called for the reference genome 1.58 were aligned against the reference genome for validation, which showed possible issues in read alignment and/or the local assembly quality. Hence, gene deletions and duplications among *C. geophilum* strains were only retained if the reference strain showed a normalised read depth between 0.7–1.3 for the same region. A gene was considered affected by a duplication or deletion if more than 50% of the coding sequence length was overlapping with a CNV region using the R package ggplot2 v. 3.3.0 (Wickham, 2011). One CNV region may contain several duplicated or deleted genes. Then, genomic distances between all genes and their closest transposable elements (TEs), and between CNV genes and their closest TEs were calculated with the *closest* function in BEDTools v 2.28.0 (Quinlan and Hall, 2010). Significance between median values was calculated using a two-sided Wilcoxon test.

### Within-species genetic diversity and signatures of selection

To improve our understanding of genomic signatures of selection, we analysed three species-specific estimates of gene diversity and their deviations from genome-wide average; nucleotide diversity per site (π), Tajima’s *D*, and *p*N/*p*S ratios. We assessed these population genetic statistics for individual genes using the R package PopGenome v. 2.6.1 (Pfeifer *et al*., 2014) and included all SNPs falling within coding sequence boundaries that were genotyped in at least 15 out of 16 *C. geophilum* strains. At the genome-wide level, a negative Tajima’s *D* is indicative of a population size expansion (e.g., after a bottleneck), whereas a positive Tajima’s *D* is supporting a decrease in population size. At the gene-specific level, negative Tajima’s *D* values are indicating of recent positive selection, while positive values are imputable to balancing selection (Hohenlohe *et al*., 2010). For π estimates, a decrease in gene-specific values in comparison to the average genome-wide level suggests a signature of positive selection (Hohenlohe *et al*., 2010). Regarding *p*N/*p*S, a ratio greater than one implies positive selection, whereas a value less than one assumes purifying selection (Tanaka and Nei, 1989). The three genotype datasets (see above) were analysed to evaluate the robustness of the results. To further examine the genes putatively under selection, we identified top-candidate genes based on the 20 lowest Tajima’s *D* values in each of the two datasets, the ref. and non-ref. cluster. Functional annotation of the top-candidate genes was carried as outlined in Peter *et al*., (2016) and according to the JGI annotations (mycocosm.jgi.doe.gov), including predicted gene ontology (GO), EuKaryotic Orthologous Groups (KOG), protein family clarification using the InterPro database (Blum *et al*., 2021), Carbohydrate-Active enZYme (CAZY; Cantarel *et al*., 2009) and secondary metabolism genes and cluster prediction (Khaldi *et al*., 2010). Gene expression data from free-living mycelium and ECM root-tips were taken from the study of Peter *et al*. 2016 and presented here. In addition, we examined the affiliation to multigene families, their PFAM domains and family size changes according to expansion/depletion analysis as detailed in Ohm *et al*. (2012) and CAFE analyses (De Bie *et al*., 2006) using the species set outlined in Peter *et al*. 2016 and available on the JGI *C. geophilum* 1.58 portal under the MCL Clusters Run “Cenococcum + 50 genomes 03/21/14.399”.

### Chromosomal synteny analysis of the MAT1 locus

We analysed chromosomal synteny of the *MAT1* locus containing the *MAT1-1* and *MAT1-2* idiomorphs based on our *de novo* genome assemblies of individual *C. geophilum* isolates. We used SPAdes v. 3.11.1 to assemble genomic scaffolds based on the paired-end Illumina sequencing data described above (Bankevich *et al*., 2012). The SPAdes pipeline includes the BayesHammer read error correction module and builds contigs in a stepwise procedure based on increasing kmer lengths. We defined a kmer range of 21, 33, 45, 67 and 89 and enabled the --careful option to reduce mismatches and indel errors in the assembly. Assembled scaffolds were error-corrected based on read realignments. We used the tool QUAST v. 4.6.0 to retrieve assembly statistics for each strain (Gurevich *et al*., 2013) and performed a blastn to identify assembled scaffolds that spanned the *MAT1* loci of the reference genome strain 1.58 (Camacho *et al*., 2009; Peter *et al*., 2016). For this, three genes in each direction of the *MAT1* locus were inspected. Once matching scaffolds were identified in individual genome assemblies, we performed pairwise blastn analyses between these scaffolds. Segmental blast hits between pairs of scaffolds known to contain the *MAT1* locus were leniently filtered for a minimum identity of 50% and a minimum alignment length of 50 bp to assess synteny patterns. For the comparison of *G. stellatum*, *L. palustris* and *C. geophilum*, we performed a tblastx search translated nucleotide and filtered additionally for a minimum bitscore of 100. We visualised pairwise synteny among scaffolds using the R package GenoPlotR v. 0.8.7 (Guy *et al*., 2011).

## Acknowledgements

We thank Stephanie Pfister and Barbara Meier for help in the laboratory; the staff at the Joint Genome Initiative for sequencing. We thank Claire Veneault-Fourrey and Joseph W. Spatafora for helpful comments and discussions. This project was supported within the framework of the ARBRE/WSL project “Blacksecret” by grants from the WSL and the French National Agency of Research (ANR), and as part of the “Investissement d’Avenir” program (ANR-11_LABX-0002-01) of Labex ARBRE (CFP15; to AK and MP) and the EC-supported Network of Excellence Evoltree (GOCE-016322 to MP).

## Data Sharing and Data Availability

Raw sequence data used in this study are available in NCBI under the Sequence Read Archive identifiers listed in Table S1.

## References

Andrews, S. (2017) FastQC: a quality control tool for high throughput sequence data.

Van der Auwera, G.A., Carneiro, M.O., Hartl, C., Poplin, R., del Angel, G., Levy-Moonshine, A., et al. (2013) From FastQ data to high-confidence variant calls: The genome analysis toolkit best practices pipeline. Curr Protoc Bioinforma 43: 1–33.

Bankevich, A., Nurk, S., Antipov, D., Gurevich, A.A., Dvorkin, M., Kulikov, A.S., et al. (2012) SPAdes: A new genome assembly algorithm and its applications to single-cell sequencing. J Comput Biol 19: 455–477.

Bazzicalupo, A.L., Ruytinx, J., Ke, Y.H., Coninx, L., Colpaert, J. V., Nguyen, N.H., et al. (2020) Fungal heavy metal adaptation through single nucleotide polymorphisms and copy-number variation. Mol Ecol 29: 4157–4169.

Berbee, M.L., Pirseyedi, M., and Hubbard, S. (1999) Cochliobolus phylogenetics and the origin of known, highly virulent pathogens, inferred from ITS and glyceraldehyde-3-phosphate dehydrogenase gene sequences. Mycologia 91: 964–977.

De Bie, T., Cristianini, N., Demuth, J.P., and Hahn, M.W. (2006) CAFE: A computational tool for the study of gene family evolution. Bioinformatics 22: 1269–1271.

Blum, M., Chang, H.Y., Chuguransky, S., Grego, T., Kandasaamy, S., Mitchell, A., et al. (2021) The InterPro protein families and domains database: 20 years on. Nucleic Acids Res 49: D344–D354.

Bolger, A.M., Lohse, M., and Usadel, B. (2014) Trimmomatic: A flexible trimmer for Illumina sequence data. Bioinformatics 30: 2114–2120.

Bourne, E.C., Mina, D., Gonçalves, S.C., Loureiro, J., Freitas, H., and Muller, L.A. (2014) Large and variable genome size unrelated to serpentine adaptation but supportive of cryptic sexuality in *Cenococcum geophilum*. Mycorrhiza 24: 13–20.

van den Brink, J. and de Vries, R.P. (2011) Fungal enzyme sets for plant polysaccharide degradation. Appl Microbiol Biotechnol 91: 1477–1492.

Bruen, T.C., Philippe, H., and Bryant, D. (2006) A simple and robust statistical test for detecting the presence of recombination. Genetics 172: 2665–2681.

Cairney, J.W. (1999) Intraspecific physiological variation: Implications for understanding functional diversity in ectomycorrhizal fungi. Mycorrhiza 9: 125–135.

Camacho, C., Coulouris, G., Avagyan, V., Ma, N., Papadopoulos, J., Bealer, K., and Madden, T.L. (2009) BLAST+: Architecture and applications. BMC Bioinformatics 10: 1–9.

Cantarel, B.I., Coutinho, P.M., Rancurel, C., Bernard, T., Lombard, V., and Henrissat, B. (2009) The Carbohydrate-Active EnZymes database (CAZy): An expert resource for glycogenomics. Nucleic Acids Res 37: 233–238.

Castresana, J. (2000) Selection of conserved blocks from multiple alignments for their use in phylogenetic analysis. Mol Biol Evol 17: 540–552.

Charlesworth, D. and Willis, J.H. (2009) The genetics of inbreeding depression. Nat Rev Genet 10: 783–796.

Chen, E.C.H., Morin, E., Beaudet, D., Noel, J., Yildirir, G., Ndikumana, S., et al. (2018) High intraspecific genome diversity in the model arbuscular mycorrhizal symbiont *Rhizophagus irregularis*. New Phytol 220: 1161–1171.

Cingolani, P., Platts, A., Wang, L.L., Coon, M., Nguyen, T., Wang, L., et al. (2012) A program for annotating and predicting the effects of single nucleotide polymorphisms, SnpEff: SNPs in the genome of Drosophila melanogaster strain w1118; iso-2; iso-3. Fly 6: 80–92.

Coleman, M.D., Bledsoe, C.S., and Lopushinsky, W. (1989) Pure culture response of ectomycorrhizal fungi to imposed water stress. Can J Bot 67: 29–39.

Daboussi, M.J. and Capy, P. (2003) Transposable elements in filamentous fungi. Annu Rev Microbiol 57: 275–299.

Danecek, P., Auton, A., Abecasis, G., Albers, C.A., Banks, E., DePristo, M.A., et al. (2011) The variant call format and VCFtools. Bioinformatics 27: 2156–2158.

Darriba, D., Posada, D., Kozlov, A.M., Stamatakis, A., Morel, B., and Flouri, T. (2020) ModelTest-NG: A new and scalable tool for the selection of DNA and protein evolutionary models. Mol Biol Evol 37: 291–294.

Debuchy, R., Berteaux-Lecellier, V., and Silar, P. (2014) Mating systems and sexual morphogenesis in ascomycetes. In Cellular and molecular biology of filamentous fungi. Borkovich, K.A. and Ebbole, D.J. (eds). Washington, U.S.A.: ASM Press, pp. 499–535.

Derbyshire, M.C. (2020) Bioinformatic detection of positive selection pressure in plant pathogens: The neutral theory of molecular sequence evolution in action. Front Microbiol 11: 1–14.

Douhan, G.W., Martin, D.P., and Rizzo, D.M. (2007) Using the putative asexual fungus *Cenococcum geophilum* as a model to test how species concepts influence recombination analyses using sequence data from multiple loci. Curr Genet 52: 191– 201.

Douhan, G.W. and Rizzo, D.M. (2005) Phylogenetic divergence in a local population of the ectomycorrhizal fungus *Cenococcum geophilum*. New Phytol 166: 263–271.

Dyrka, W., Lamacchia, M., Durrens, P., Kobe, B., Daskalov, A., Paoletti, M., et al. (2014) Diversity and variability of NOD-like receptors in fungi. Genome Biol Evol 6: 3137–3158.

Edler, D., Klein, J., Antonelli, A., and Silvestro, D. (2021) raxmlGUI 2.0: A graphical interface and toolkit for phylogenetic analyses using RAxML. Methods Ecol Evol 12: 373–377.

Falush, D., Stephens, M., and Pritchard, J.K. (2003) Inference of population structure using multilocus genotype data: Linked loci and correlated allele frequencies. Genetics 164: 1567–1587.

Fernandez, C.W., McCormack, M.L., Hill, J.M., Pritchard, S.G., and Koide, R.T. (2013) On the persistence of *Cenococcum geophilum* ectomycorrhizas and its implications for forest carbon and nutrient cycles. Soil Biol Biochem 65: 141–143.

de Freitas Pereira, M., Veneault-Fourrey, C., Vion, P., Guinet, F., Morin, E., Barry, K.W., et al. (2018) Secretome analysis from the ectomycorrhizal ascomycete *Cenococcum geophilum*. Front Microbiol 9: 1–17.

Gardes, M. and Bruns, T.D. (1993) ITS primers with enhanced specificity for basidiomycetes - application to the identification of mycorrhizae and rusts. Mol Ecol 2: 113–118.

Geml, J., Laursen, G.A., O’Neill, K., Nusbaum, H.C., and Taylor, D.L. (2006) Beringian origins and cryptic speciation events in the fly agaric (*Amanita muscaria*). Mol Ecol 15: 225–239.

Gioti, A., Mushegian, A.A., Strandberg, R., Stajich, J.E., and Johannesson, H. (2012) Unidirectional evolutionary transitions in fungal mating systems and the role of transposable elements. Mol Biol Evol 29: 3215–3226.

Gonçalves, S.C., Martins-Loução, M.A., and Freitas, H. (2009) Evidence of adaptive tolerance to nickel in isolates of *Cenococcum geophilum* from serpentine soils. Mycorrhiza 19: 221–230.

Gurevich, A., Saveliev, V., Vyahhi, N., and Tesler, G. (2013) QUAST: Quality assessment tool for genome assemblies. Bioinformatics 29: 1072–1075.

Guy, L., Kultima, J.R., Andersson, S.G.E., and Quackenbush, J. (2011) genoPlotR: comparative gene and genome visualization in R. Bioinformatics 27: 2334–2335.

Hartmann, F.E. and Croll, D. (2017) Distinct trajectories of massive recent gene gains and losses in populations of a microbial eukaryotic pathogen. Mol Biol Evol 34: 2808–2822.

Hartmann, F.E., McDonald, B.A., and Croll, D. (2018) Genome-wide evidence for divergent selection between populations of a major agricultural pathogen. Mol Ecol 27: 2725– 2741.

Hasselquist, N., Germino, M.J., McGonigle, T., and Smith, W.K. (2005) Variability of *Cenococcum* colonization and its ecophysiological significance for young conifers at alpine-treeline. New Phytol 165: 867–873.

Hemsworth, G.R., Henrissat, B., Davies, G.J., and Walton, P.H. (2014) Discovery and characterization of a new family of lytic polysaccharide monooxygenases. Nat Chem Biol 10: 122–126.

Herzog, C., Peter, M., Pritsch, K., Günthardt-Goerg, M.S., and Egli, S. (2013) Drought and air warming affects abundance and exoenzyme profiles of *Cenococcum geophilum* associated with *Quercus robur*, Q. petraea and Q. pubescens. Plant Biol 15: 230–237.

Hohenlohe, P.A., Phillips, P.C., and Cresko, W.A. (2010) Using population genomics to detect selection in natural populations: Key concepts and methodological considerations. Int J Plant Sci 171: 1059–1071.

Huson, D.H. and Bryant, D. (2006) Application of phylogenetic networks in evolutionary studies. Mol Biol Evol 23: 254–267.

Johnson, D., Martin, F., Cairney, J.W., and Anderson, I.C. (2012) The importance of individuals: Intraspecific diversity of mycorrhizal plants and fungi in ecosystems. New Phytol 194: 614–628.

Katoh, K., Rozewicki, J., and Yamada, K.D. (2019) MAFFT online service: Multiple sequence alignment, interactive sequence choice and visualization. Brief Bioinform 20: 1160–1166.

Kerner, R., Delgado-Eckert, E., Del Castillo, E., Müller-Starck, G., Peter, M., Kuster, B., et al. (2012) Comprehensive proteome analysis in *Cenococcum geophilum* Fr. as a tool to discover drought-related proteins. J Proteomics 75: 3707–3719.

Khaldi, N., Seifuddin, F.T., Turner, G., Haft, D., Nierman, W.C., Wolfe, K.H., and Fedorova, N.D. (2010) SMURF: Genomic mapping of fungal secondary metabolite clusters. Fungal Genet Biol 47: 736–741.

Kozlov, A.M., Darriba, D., Flouri, T., Morel, B., and Stamatakis, A. (2019) RAxML-NG: A fast, scalable and user-friendly tool for maximum likelihood phylogenetic inference. Bioinformatics 35: 4453–4455.

Langmead, B., Trapnell, C., Pop, M., and Salzberg, S.L. (2009) Ultrafast and memory-efficient alignment of short DNA sequences to the human genome. Genome Biol 10: 1– 10.

Larsen, P.E., Sreedasyam, A., Trivedi, G., Desai, S., Dai, Y., Cseke, L.J., and Collart, F.R. (2016) Multi-omics approach identifies molecular mechanisms of plant-fungus mycorrhizal interaction. Front Plant Sci 6: 1–17.

Li, H., Handsaker, B., Wysoker, A., Fennell, T., Ruan, J., Homer, N., et al. (2009) The sequence alignment/map format and SAMtools. Bioinformatics 25: 2078–2079.

Li, W., Sullivan, T.D., Walton, E., Averette, A.F., Sakthikumar, S., Cuomo, C.A., et al. (2013) Identification of the mating-type (*MAT*) locus that controls sexual reproduction of blastomyces dermatitidis. Eukaryot Cell 12: 109–117.

van der Linde, S., Suz, L.M., Orme, D. l., Cox, F., Andreae, H., Asi, E., et al. (2018) Environment and host as large-scale controls of ectomycorrhizal fungi. Nature 558: 243–248.

LoBuglio, K.F. (1999) Cenococcum. In Ectomycorrhizal Fungi: Key Genera in Profile. Cairney, J.W.G. and Chambers, S.M. (eds). Heidelberg, Germany: Springer, pp. 287–309.

LoBuglio, K.F. and Taylor, J.W. (2002) Recombination and genetic differentiation in the mycorrhizal fungus *Cenococcum geophilum* Fr. Mycologia 94: 772–780.

Lofgren, L.A., Nguyen, N.H., Vilgalys, R., Ruytinx, J., Liao, H.L., Branco, S., et al. (2021) Comparative genomics reveals dynamic genome evolution in host specialist ectomycorrhizal fungi. New Phytol 230: 774–792.

Martin, F., Kohler, A., Murat, C., Balestrini, R., Coutinho, P.M., Jaillon, O., et al. (2010) Périgord black truffle genome uncovers evolutionary origins and mechanisms of symbiosis. Nature 464: 1033–1038.

Martin, F., Kohler, A., Murat, C., Veneault-Fourrey, C., and Hibbett, D.S. (2016) Unearthing the roots of ectomycorrhizal symbioses. Nat Rev Microbiol 14: 760–773.

McKenna, A., Hanna, M., Banks, E., Sivachenko, A.Y., Cibulskis, K., Kernytsky, A.M., et al. (2010) The genome analysis toolkit: A MapReduce framework for analyzing next-generation DNA sequencing data. Genome Res 20: 1297–1303.

Miyauchi, S., Kiss, E., Kuo, A., Drula, E., Kohler, A., Sánchez-García, M., et al. (2020) Large-scale genome sequencing of mycorrhizal fungi provides insights into the early evolution of symbiotic traits. Nat Commun 11: 1–17.

Mujic, A.B., Durall, D.M., Spatafora, J.W., and Kennedy, P.G. (2016) Competitive avoidance not edaphic specialization drives vertical niche partitioning among sister species of ectomycorrhizal fungi. New Phytol 209: 1174–1183.

Muszewska, A., Steczkiewicz, K., Stepniewska-Dziubinska, M., and Ginalski, K. (2019) Transposable elements contribute to fungal genes and impact fungal lifestyle. Sci Rep 9: 1–10.

Nygren, K., Strandberg, R., Wallberg, A., Nabholz, B., Gustafsson, T., García, D., et al. (2011) A comprehensive phylogeny of *Neurospora* reveals a link between reproductive mode and molecular evolution in fungi. Mol Phylogenet Evol 59: 649–663.

Obase, K., Douhan, G.W., Matsuda, Y., and Smith, M.E. (2014) Culturable fungal assemblages growing within *Cenococcum* sclerotia in forest soils. FEMS Microbiol Ecol 90: 708–717.

Obase, K., Douhan, G.W., Matsuda, Y., and Smith, M.E. (2018) Isolation source matters: Sclerotia and ectomycorrhizal roots provide different views of genetic diversity in *Cenococcum geophilum*. Mycologia 110: 473–481.

Obase, K., Douhan, G.W., Matsuda, Y., and Smith, M.E. (2017) Progress and challenges in understanding the biology, diversity, and biogeography of *Cenococcum geophilum*. In Biogeography of mycorrhizal symbiosis. Tedersoo, L. (ed)., pp. 299–317.

Obase, K., Douhan, G.W., Matsuda, Y., and Smith, M.E. (2016) Revisiting phylogenetic diversity and cryptic species of *Cenococcum geophilum* sensu lato. Mycorrhiza 26: 529– 540.

Ohm, R.A., Feau, N., Henrissat, B., Schoch, C.L., Horwitz, B.A., Barry, K.W., et al. (2012) Diverse lifestyles and strategies of plant pathogenesis encoded in the genomes of eighteen Dothideomycetes fungi. PLoS Pathog 8:.

Paoletti, M. (2016) Vegetative incompatibility in fungi: From recognition to cell death, whatever does the trick. Fungal Biol Rev 30: 152–162.

Payen, T., Murat, C., Gigant, A., Morin, E., De Mita, S., and Martin, F. (2015) A survey of genome-wide single nucleotide polymorphisms through genome resequencing in the Périgord black truffle (*Tuber melanosporum* Vittad.). Mol Ecol Resour 15: 1243–1255.

Peter, M., Kohler, A., Ohm, R.A., Kuo, A., Krützmann, J., Morin, E., et al. (2016) Ectomycorrhizal ecology is imprinted in the genome of the dominant symbiotic fungus *Cenococcum geophilum*. Nat Commun 7: 12662.

Pfeifer, B., Wittelsbürger, U., Ramos-Onsins, S.E., and Lercher, M.J. (2014) PopGenome: An efficient Swiss army knife for population genomic analyses in R. Mol Biol Evol 31: 1929–1936.

di Pietro, M., Churin, J.L., and Garbaye, J. (2007) Differential ability of ectomycorrhizas to survive drying. Mycorrhiza 17: 547–550.

Pritchard, J.K., Stephens, M., and Donnelly, P. (2000) Inference of population structure using multilocus genotype data. Genetics 155: 945–959.

Quinlan, A.R. and Hall, I.M. (2010) BEDTools: A flexible suite of utilities for comparing genomic features. Bioinformatics 26: 841–842.

Raffaele, S. and Kamoun, S. (2012) Genome evolution in filamentous plant pathogens: Why bigger can be better. Nat Rev Microbiol 10: 417–430.

Reinhardt, D., Roux, C., Corradi, N., and Di Pietro, A. (2021) Lineage-specific genes and cryptic sex: Parallels and differences between arbuscular mycorrhizal fungi and fungal pathogens. Trends Plant Sci 26: 111–123.

Riley, R. and Corradi, N. (2013) Searching for clues of sexual reproduction in the genomes of arbuscular mycorrhizal fungi. Fungal Ecol 6: 44–49.

Rineau, F. and Courty, P.E. (2011) Secreted enzymatic activities of ectomycorrhizal fungi as a case study of functional diversity and functional redundancy. Ann For Sci 68: 69–80.

Rochet, J., Moreau, P.A., Manzi, S., and Gardes, M. (2011) Comparative phylogenies and host specialization in the alder ectomycorrhizal fungi Alnicola, Alpova and Lactarius (Basidiomycota) in Europe. BMC Evol Biol 11: 1–14.

Sánchez-Ramírez, S., Tulloss, R.E., Guzmán-Dávalos, L., Cifuentes-Blanco, J., Valenzuela, R., Estrada-Torres, A., et al. (2015) In and out of refugia: Historical patterns of diversity and demography in the North American Caesar’s mushroom species complex. Mol Ecol 24: 5938–5956.

Serrato-Capuchina, A. and Matute, D.R. (2018) The role of transposable elements in speciation. Genes 9: 1–29.

Simpson, M.C., Coetzee, M.P.A., van der Nest, M.A., Wingfield, M.J., and Wingfield, B.D. (2018) Ceratocystidaceae exhibit high levels of recombination at the mating-type (*MAT*) locus. Fungal Biol 122: 1184–1191.

Smith, M.E., Henkel, T.W., and Rollins, J.A. (2015) How many fungi make sclerotia? Fungal Ecol 13: 211–220.

Spatafora, J.W., Owensby, C.A., Douhan, G.W., Boehm, E.W.A., and Schoch, C.L. (2012) Phylogenetic placement of the ectomycorrhizal genus *Cenococcum* in Gloniaceae (Dothideomycetes). Mycologia 104: 758–765.

Tanaka, T. and Nei, M. (1989) Positive Darwinian selection observed at the variable-region genes of immunoglobulins. Mol Biol Evol 6: 447–459.

Trappe, J.M. (1962) Cenococcum graniforme - Its distribution, ecology, mycorrhiza formation, and inherent variation.

Vélez, J., Morris, R., Vilgalys, R., Labbé, J., and Schadt, C. (2021) Phylogenetic diversity of 200+ isolates of the ectomycorrhizal fungus *Cenococcum geophilum* associated with *Populus trichocarpa* soils in the Pacific Northwest, USA and comparison to globally distributed representatives. PLoS One 16: e0231367.

Wang, N.Y., Zhang, K., Huguet-Tapia, J.C., Rollins, J.A., and Dewdney, M.M. (2016) Mating type and simple sequence repeat markers indicate a clonal population of *Phyllosticta citricarpa* in Florida. Phytopathology 106: 1300–1310.

Wang, X., Zheng, Z., Cai, Y., Chen, T., Li, C., Fu, W., and Jiang, Y. (2017) CNVcaller: Highly efficient and widely applicable software for detecting copy number variations in large populations. Gigascience 6: 1–12.

Watanabe, M., Sato, H., Matsuzaki, H., Kobayashi, T., Sakagami, N., Maejima, Y., et al. (2007) 14C ages and δ13C of sclerotium grains found in forest soils. Soil Sci Plant Nutr 53: 125–131.

White, T., Bruns, T., Lee, S., and Taylor, J. (1990) Amplification and direct sequencing of fungal ribosomal RNA genes for phylogenetics. In PCR protocols: a guide to methods and applications. pp. 315–322.

Wickham, H. (2011) ggplot2. Wiley Interdiscip Rev Comput Stat 3: 180–185.

Zeiner, C.A., Purvine, S.O., Zink, E.M., Paša-Tolić, L., Chaput, D.L., Haridas, S., et al. (2016) Comparative analysis of secretome profiles of manganese(II)-oxidizing Ascomycete fungi. PLoS One 11:.

Zhang, J., Kapli, P., Pavlidis, P., and Stamatakis, A. (2013) A general species delimitation method with applications to phylogenetic placements. Bioinformatics 29: 2869–2876.

Zheng, X., Levine, D., Shen, J., Gogarten, S.M., Laurie, C., and Weir, B.S. (2012) A high-performance computing toolset for relatedness and principal component analysis of SNP data. Bioinformatics 28: 3326–3328.

