## Supplementary Information for "Cryptic genetic structure and copy-number variation in the ubiquitous forest symbiotic fungus *Cenococcum geophilum*"

Affiliations:
[†]Swiss Federal Research Institute WSL, Birmensdorf, Switzerland
[¶]INRAE, UMR 1136 INRAE-University of Lorraine, Interactions Arbres/Microorganismes, Laboratory of Excellence ARBRE, INRAE-Grand Est, Champenoux, France
[§]Department of Plant and Microbial Biology, University of California, Berkeley, USA
[#]U.S. Department of Energy Joint Genome Institute, Lawrence Berkeley National Laboratory, Berkeley, USA
[‡]Laboratory of Evolutionary Genetics, Institute of Biology, University of Neuchâtel, Neuchâtel, Switzerland

[*]Co-last authors contributed equally to this study

Tables S1–S4 and Figs. S1–S4

**Table S1 Summary of the genome sequencing data with mapping statistics from the studied *Cenococcum geophilum* strains.**

| Strain | BioProject ID | Sequencing ID | SRA ID | SRA ID | Total raw reads | Overall alignment percent |
|---|---|---|---|---|---|---|
| 1.58 | PRJNA196023 | SRR1800544 | SRS2328413 | SRP003514 | 45,519,993 | 99.29 |
| 1.094 | PRJNA334340 | SRR4136285 | SRS1667730 | SRP084857 | 6,737,436 | 71.28 |
| 1.094 | PRJNA334340 | SRR4136286 | SRS1667730 | SRP084857 | 6,261,388 | 71.31 |
| 1.075 | PRJNA334338 | SRR4136288 | SRS1667732 | SRP084859 | 11,332,369 | 77.56 |
| 1.075 | PRJNA334338 | SRR4136289 | SRS1667732 | SRP084859 | 10,404,315 | 78.10 |
| 1.096 | PRJNA334341 | SRR4136290 | SRS1667733 | SRP084860 | 10,700,440 | 64.37 |
| 1.096 | PRJNA334341 | SRR4136291 | SRS1667733 | SRP084860 | 9,884,876 | 64.80 |
| 1.104 | PRJNA334346 | SRR4136292 | SRS1667734 | SRP084861 | 10,218,188 | 90.10 |
| 1.104 | PRJNA334346 | SRR4136293 | SRS1667734 | SRP084861 | 9,618,457 | 90.15 |
| 1.076 | PRJNA334339 | SRR4136296 | SRS1667736 | SRP084863 | 11,093,698 | 94.16 |
| 1.076 | PRJNA334339 | SRR4136297 | SRS1667736 | SRP084863 | 10,325,455 | 94.21 |
| 1.022 | PRJNA334333 | SRR4136298 | SRS1667737 | SRP084864 | 11,619,059 | 80.40 |
| 1.022 | PRJNA334333 | SRR4136299 | SRS1667737 | SRP084864 | 10,865,314 | 80.87 |
| 1.100 | PRJNA334343 | SRR4136300 | SRS1667738 | SRP084865 | 12,078,625 | 78.03 |
| 1.100 | PRJNA334343 | SRR4136301 | SRS1667738 | SRP084865 | 11,216,107 | 78.55 |
| 1.061 | PRJNA334336 | SRR4136302 | SRS1667739 | SRP084866 | 9,337,691 | 95.43 |
| 1.061 | PRJNA334336 | SRR4136303 | SRS1667739 | SRP084866 | 8,719,813 | 95.54 |
| 1.097 | PRJNA334342 | SRR4136306 | SRS1667741 | SRP084868 | 10,292,074 | 90.69 |
| 1.097 | PRJNA334342 | SRR4136307 | SRS1667741 | SRP084868 | 9,547,525 | 90.71 |
| 1.101 | PRJNA334344 | SRR4136308 | SRS1667742 | SRP084869 | 12,396,544 | 81.88 |
| 1.101 | PRJNA334344 | SRR4136309 | SRS1667742 | SRP084869 | 11,470,363 | 82.19 |
| 1.060 | PRJNA334335 | SRR4136310 | SRS1667743 | SRP084870 | 10,053,052 | 86.72 |
| 1.060 | PRJNA334335 | SRR4136311 | SRS1667743 | SRP084870 | 9,379,193 | 86.80 |
| 1.064 | PRJNA334337 | SRR4136312 | SRS1667744 | SRP084871 | 10,564,580 | 77.09 |
| 1.064 | PRJNA334337 | SRR4136313 | SRS1667744 | SRP084871 | 9,755,114 | 77.80 |
| 1.102 | PRJNA334345 | SRR4136314 | SRS1667745 | SRP084872 | 10,490,785 | 91.58 |
| 1.102 | PRJNA334345 | SRR4136315 | SRS1667745 | SRP084872 | 9,799,524 | 91.59 |
| 1.057 | PRJNA334334 | SRR4136316 | SRS1667746 | SRP084873 | 10,983,999 | 87.08 |
| 1.057 | PRJNA334334 | SRR4136317 | SRS1667746 | SRP084873 | 10,266,328 | 87.59 |
| 1.005 | PRJNA334332 | SRR4136320 | SRS1667748 | SRP084875 | 9,265,475 | 86.40 |
| 1.005 | PRJNA334332 | SRR4136321 | SRS1667748 | SRP084875 | 8,697,123 | 86.63 |

**Table S2 Characteristics of the two barcoded nuclear regions and substitution models used in the maximum likelihood phylogenetic tree analysis.**

| Locus | Aligned length (bp) | Variable sites (No.) | Invariant sites (%) | Gaps (%) | AIC score | AICc sore | BIC score | Model implemented in RAxML |
|---|---|---|---|---|---|---|---|---|
| GAPDH | 597 | 221 | 66.50 | 2.73 | 5402.5958 | 5429.5958 | 5767.1249 | SYM + G |
| ITS | 949 | 277 | 78.71 | 13.60 | 7001.4655 | 7015.4655 | 7389.8982 | K80 + I + G |
| Two loci partitioned | 1546 | 498 | 74.00 | 9.40 | – | – | – | SYM + G, K80 + I + G |

**Table S3 Recombination analysis from the *Cenococcum geophilum* strains analysed in this study.**

| Dataset | Mean | Variance | Observed | *P* value |
|---|---|---|---|---|
| All | 0.271 | $0.529 \times 10^{-7}$ | 0.262 | < 0.001 |
| Ref. cluster | 0.383 | $3.827 \times 10^{-7}$ | 0.356 | < 0.001 |
| Non-ref. cluster | 0.166 | $2.522 \times 10^{-7}$ | 0.166 | 0.641 |

**Table S4 Annotation of top-candidate genes.**

See separate file.

**Table S5 Characteristics of the *Cenococcum geophilum de novo* genome assemblies analysed in this study.**

| Strain | Number of contigs | Largest contig (bp) | Total length (bp) | N50 | L50 | GC content (%) | Mismatches per 100 kbp |
|---|---|---|---|---|---|---|---|
| 1.005 | 22,839 | 190,928 | 132,419,543 | 12,598 | 2,439 | 37.55 | 113.43 |
| 1.022 | 37,018 | 132,204 | 110,597,253 | 5,720 | 3,460 | 39.62 | 171.64 |
| 1.075 | 27,259 | 153,375 | 164,410,048 | 11,927 | 3,431 | 36.21 | 144.15 |
| 1.076 | 37,593 | 161,367 | 179,246,920 | 9,154 | 4,759 | 36.75 | 236.90 |
| 1.094 | 14,891 | 181,770 | 182,540,467 | 23,528 | 2,247 | 32.61 | 260.41 |
| 1.096 | 14,654 | 193,835 | 188,054,450 | 24,382 | 2,255 | 32.6 | 79.41 |
| 1.101 | 36,800 | 172,864 | 110,264,397 | 5,822 | 3,440 | 39.62 | 61.49 |
| 1.102 | 28,945 | 144,554 | 166,007,074 | 10,659 | 3,941 | 37.82 | 123.78 |
| 1.104 | 22,664 | 162,548 | 129,798,722 | 13,112 | 2,404 | 38.15 | 64.07 |

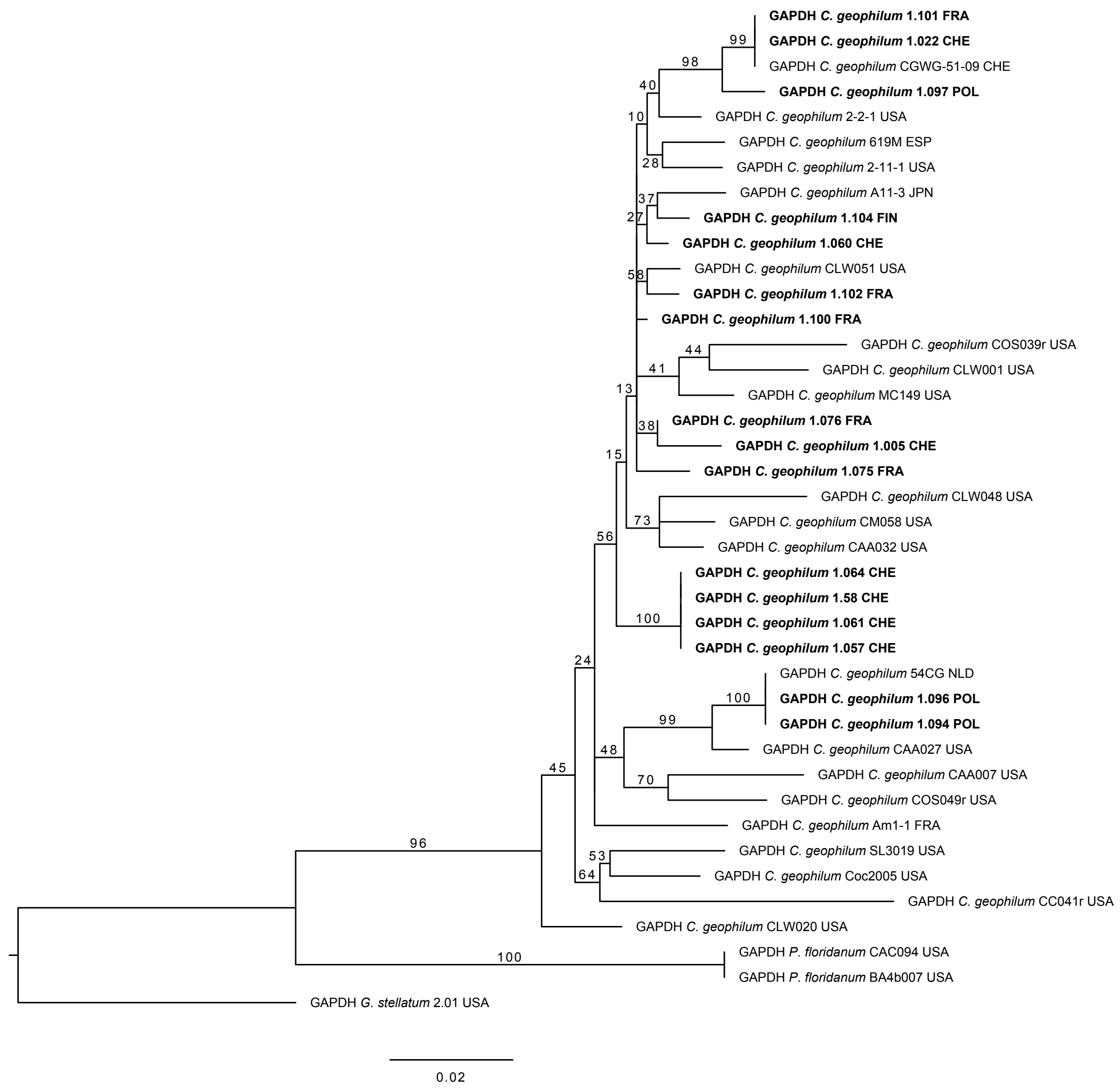


**Fig. S1 Maximum likelihood phylogenetic tree of the GAPDH region depicting the relationships between the *Cenococcum geophilum* lineages and closely related species.** Abbreviations "*C.*", "*P.*", and "*G.*" refer to the genus names *Cenococcum*, *Pseudocenococcum*, and *Glonium*, respectively. Topology was rooted with *Glonium stellatum* as outgroup. Branch supports represent bootstrap values from the maximum likelihood tree reconstruction. The scale bar shows the substitution rate. Tips in bold are the 16 strains analysed in this study using whole-genome sequencing data and the sample codes are described in Table S1.

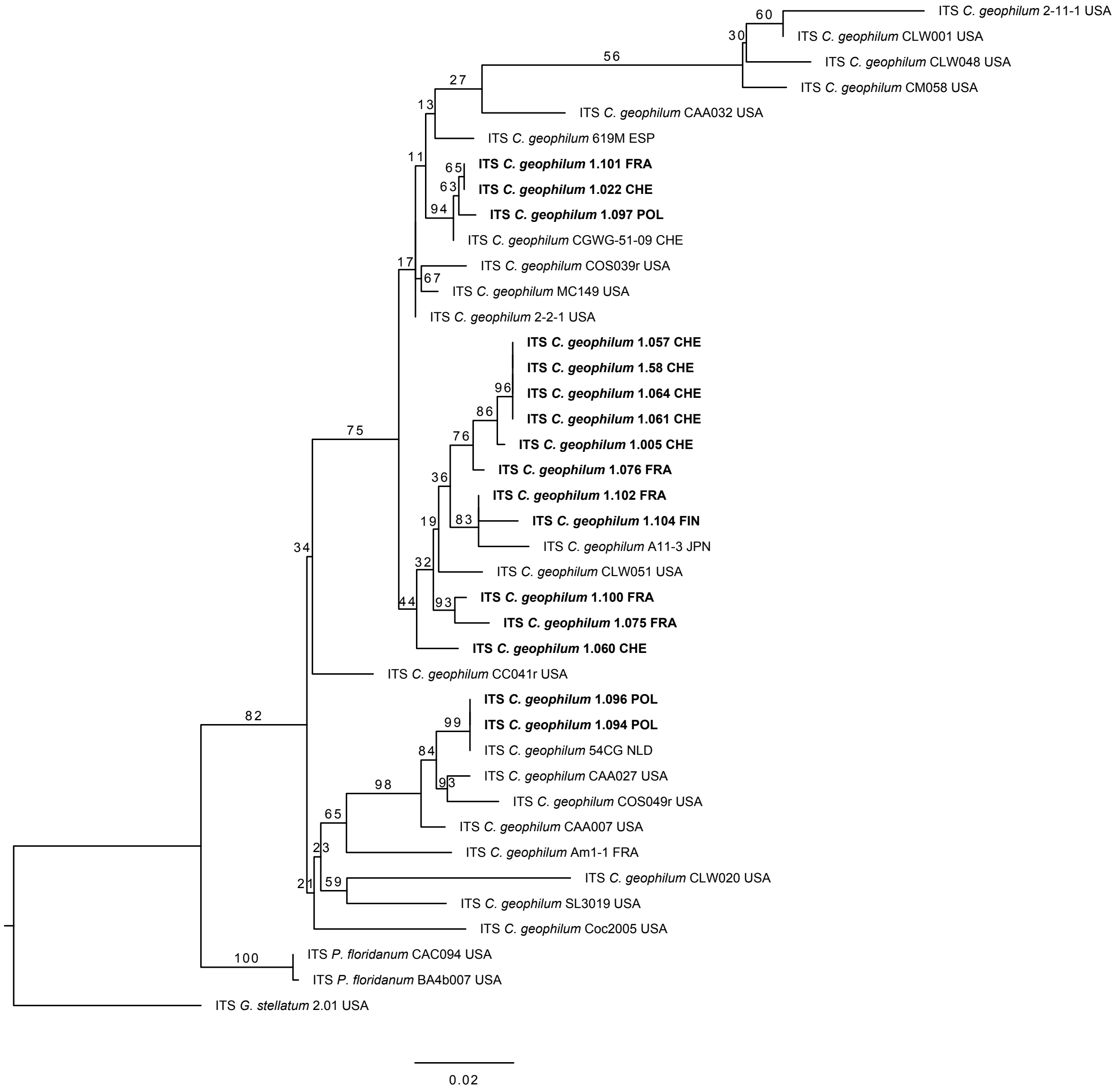


**Fig. S2 Maximum likelihood phylogenetic tree of the ITS region depicting the relationships between the *Cenococcum geophilum* lineages and closely related species.** Abbreviations "*C.*", "*P.*", and "*G.*" refer to the genus names *Cenococcum*, *Pseudocenococcum*, and *Glonium*, respectively. Topology was rooted with *Glonium stellatum* as outgroup. Branch supports represent bootstrap values from the maximum likelihood tree reconstruction. The scale bar shows the substitution rate. Tips in bold are the 16 strains analysed in this study using whole-genome sequencing data and the sample codes are described in Table S1.

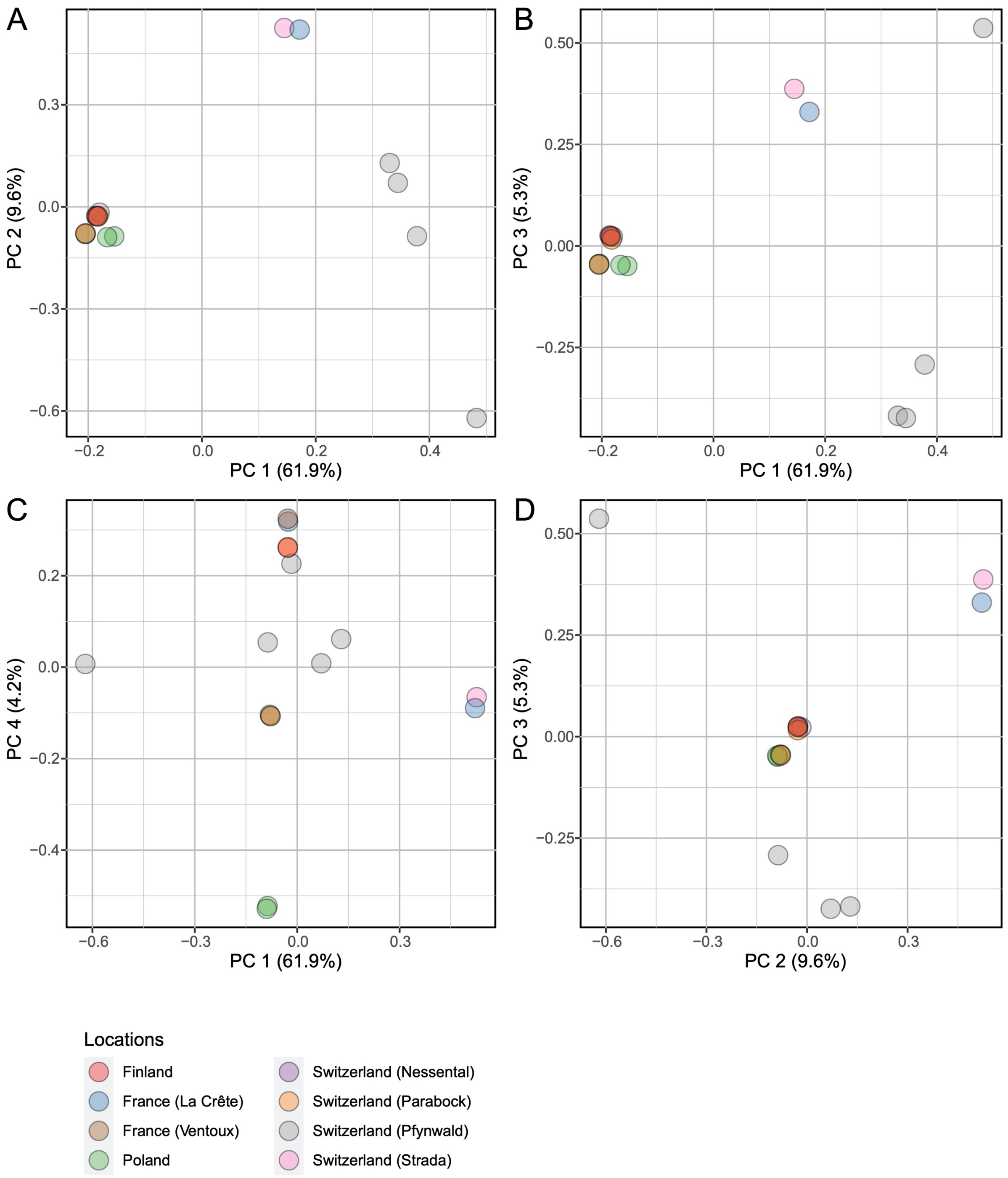


**Fig. S3 Genetic structure of *Cenococcum geophilum* individuals based on an ordination method.** Principal component (PC) analysis showing **A**, PC1 and PC2, **B**, PC1 and PC3, **C**, PC1 and PC4, and **D**, PC2 and PC3. Percent values noted in the axis labels refer to the percentage of total variance explained. Colours indicate the population of each strain. Strain codes are explained in Table S1.

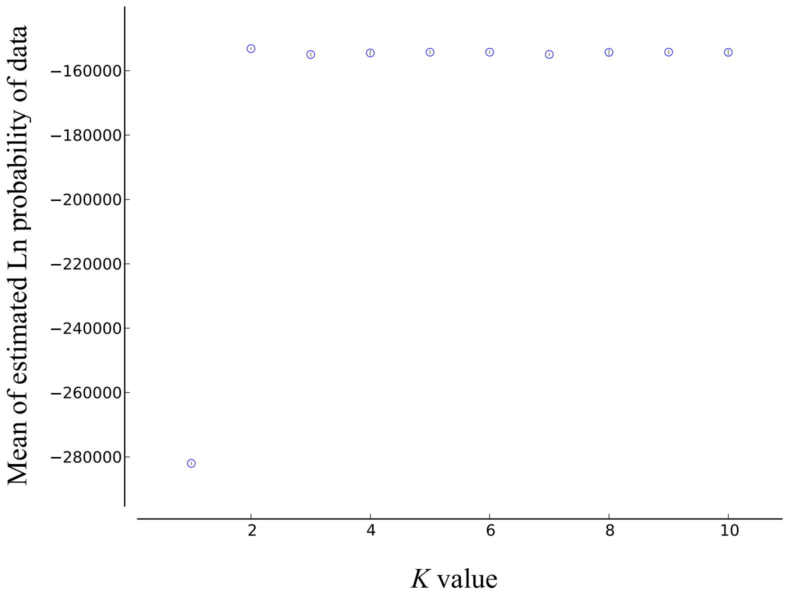


**Fig. S4 Distribution of estimated Ln probability of data across *K* values in Bayesian clustering analysis.** Mean and standard deviation of each *K* value were assessed from five replications.
